# The interferon-inducible GTPase MxB promotes capsid disassembly and genome release of herpesviruses

**DOI:** 10.1101/2022.01.25.477704

**Authors:** Manutea C. Serrero, Virginie Girault, Sebastian Weigang, Todd M. Greco, Ana Ramos-Nascimento, Fenja Anderson, Antonio Piras, Ana Hickford Martinez, Jonny Hertzog, Anne Binz, Anja Pohlmann, Ute Prank, Jan Rehwinkel, Rudolf Bauerfeind, Ileana M. Cristea, Andreas Pichlmair, Georg Kochs, Beate Sodeik

## Abstract

Host proteins sense viral products and induce defence mechanisms, particularly in immune cells. Using cell-free assays and quantitative mass spectrometry, we determined the interactome of capsid-host protein complexes of herpes simplex virus and identified the large dynamin-like GTPase myxovirus resistance protein B (MxB) as an interferon-inducible protein interacting with capsids. Electron microscopy analyses showed that cytosols containing MxB had the remarkable capability to disassemble the icosahedral capsids of herpes simplex viruses and varicella zoster virus into flat sheets of connected triangular faces. In contrast, capsids remained intact in cytosols with MxB mutants unable to hydrolyse GTP or to dimerize. Our data suggest that MxB senses herpesviral capsids, mediates their disassembly, and thereby restricts the efficiency of nuclear targeting of incoming capsids and/or the assembly of progeny capsids. The resulting premature release of viral genomes from capsids may enhance the activation of DNA sensors, and thereby amplify the innate immune responses.

## INTRODUCTION

Infections with human alphaherpesviruses are associated with painful and stigmatizing manifestations such as herpes labialis or herpes genitalis, but also cause life-threatening meningitis or encephalitis, potentially blinding eye infections, herpes zoster, and post-herpetic neuralgia, particularly in immunocompromised patients (Gershon *et al*., 2015; Whitley & Roizman, 2016; Whitley & Johnston, 2021). Herpes simplex viruses (HSV-1, HSV-2) and varicella zoster virus (VZV) productively infect epithelial and fibroblast cells of the skin and mucous membranes as well as neurons, but are restricted in immune cells. Macrophages, Langerhans cells, dendritic cells, and NK cells mount potent immune responses against alphaherpesviruses (Whitley & Roizman, 2016).

Intracellular DNA sensors are crucial to sense herpesvirus infections, and to induce caspase-1 mediated inflammation and type I IFN expression (Hertzog & Rehwinkel, 2020; Kurt-Jones *et al*., 2017; Lum & Cristea, 2021, Ma *et al*., 2018; Paludan *et al*., 2019; Stempel *et al*., 2019). During an unperturbed infection, capsid shells shield herpesviral genomes from cytosolic sensors during nuclear targeting as well as after nuclear genome packaging (Arvin & Abendroth, 2021; Döhner *et al*., 2021; Knipe *et al*., 2021). HSV-1 capsids can withstand compressive forces of up to 6 nN which is more than sufficient to endure the 18 atm repulsive pressure of the packaged viral DNA (Bauer *et al*., 2013; Roos *et al*., 2009). So far, it is unclear how cytosolic DNA sensors gain access to herpesviral genomes; either cytosolic host factors disassemble the sturdy herpesviral capsids during infection, or the nuclear envelopes become leaky.

HSV-1 virions contain an amorphous tegument layer that links the icosahedral capsids with a diameter of 125 nm to the viral envelope proteins (Crump, 2018; Dai & Zhou, 2018; Diefenbach, 2015). To identify cytosolic proteins that promote or restrict infection by interacting with HSV-1 capsids, we have developed cell-free methods to reconstitute capsid-host protein complexes using tegumented capsids from extracellular viral particles or tegument-free capsids from the nuclei of infected cells (Radtke *et al*., 2014). Intact capsids are incubated with cytosol prepared from tissues or cultured cells, and the capsid-host protein complexes are isolated, and characterized by mass spectrometry (MS), immunoblot, electron microscopy, and functional assays. We could show that HSV-1 capsids require inner tegument proteins to recruit microtubule motors, to move along microtubules, to dock at nuclear pore complexes (NPCs), to release viral genomes from capsids, and to import viral genomes into the nucleoplasm, and that capsids lacking tegument cannot move along microtubules, but still bind to nuclear pores (Anderson *et al*., 2014; Ojala *et al*., 2000; Radtke *et al*., 2010; Wolfstein *et al*., 2006).

Here, we searched for proteins that might contribute to sensing cytosolic capsids and thereby promote the detection of herpesviral genomes. Using extracts of matured THP-1 cells, a model system for human macrophages (Tsuchiya *et al*., 1980), we identified type I interferon (IFN) inducible proteins that bound specifically to HSV-1 capsids. Among them was the large dynamin-like GTPase myxovirus resistance protein B (MxB). MxB limits the infection of several herpesviruses, and can mediate almost 50% of the IFN-mediated restriction of HSV-1, although its mode of action has remained elusive so far (Crameri *et al*., 2018; Liu *et al*., 2012; Schilling *et al*., 2018; Vasudevan *et al*., 2018). MxB has been first described for its potent inhibition of HIV infection (Goujon *et al*., 2013; Kane *et al*., 2013; Liu *et al*., 2013). The human *MX2* gene codes for a full-length MxB (residues 1-715) and a smaller version (residues 26-715) that lacks an N-terminal extension (NTE), which both are highly expressed upon IFN induction (Melen *et al*., 1996). MxB likely operates as an anti-parallel dimer but can also form higher-order filaments; its N-terminal GTPase domain connects to a bundle signalling element that moves relative to the GTPase domain in response to nucleotide binding, and the C-terminal stalk domain is critical for MxB oligomerization (Alvarez *et al*., 2017; Chen *et al*., 2017; Fribourgh *et al*., 2014; Gao *et al*., 2011).

We show here that both, full-length MxB(1-715) and MxB(26-715) have the remarkable property to disassemble the capsids of the three human alphaherpesviruses HSV-1, HSV-2, and VZV, so that they can no longer transport nor shield the viral genomes. Capsid disassembly did not require proteases but depended on the ability of MxB to hydrolyse GTP and to dimerize. As the large tegument protein pUL36 links the capsid vertices to the other tegument proteins (Crump, 2018; Dai & Zhou, 2018; Diefenbach, 2015), and as an increasing amount of associated tegument proteins protected capsids against MxB mediated disassembly, we propose that MxB attacks the capsids at their vertices. Our data suggest that MxB can bind to and disassemble incoming as well as progeny capsids, and thereby might increase the sensing of cytosolic and nuclear viral genomes. Therefore, the MxB GTPase might be the sought-after capsid destroyer that acts upstream of cytosolic or nuclear sensors to promote viral genome detection and induction of innate immune responses.

## RESULTS

### IFN induction prevents HSV-1 infection of macrophages

Before investigating capsid interactions with macrophage proteins, we compared HSV-1 infection in human keratinocytes (HaCat), pigment epithelial cells (RPE), and THP-1 cells at low, moderate or high multiplicity of infection (MOI). We stimulated monocyte THP-1 cells with phorbol 12-myristate 13-acetate to differentiate them into a macrophage-like phenotype, and used them either directly (Mφ) or after a resting period of 3 days (Mφ_R_). HSV-1 replicated productively in HaCat and RPE cells up to 20 hpi, while a pre-treatment with IFN delayed and reduced but did not prevent the production of infectious virions (Fig. 1). Both Mφ and Mφ_R_ released 10 to 100-fold less infectious HSV-1, and an IFN pre-treatment prevented infection at all MOIs. Thus, Mφ and Mφ_R_ restricted HSV-1 infection efficiently, and the induction of IFN-stimulated genes (ISGs) prevented any productive infection.

**Figure 1:**
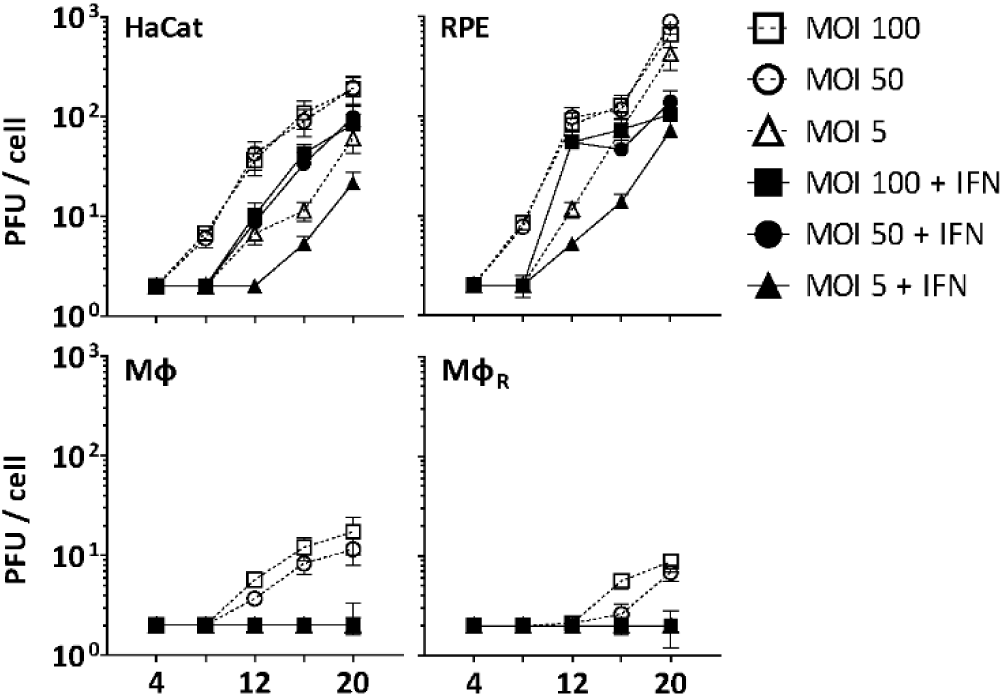
IFN restricts HSV-1 infection in keratinocytes, epithelial cells, and macrophages. HaCat, RPE, Mφ, or Mφ_R_ cells were mock-treated or treated with human IFN-α (1000 U/mL) for 16 h and were infected with HSV-1(17^+^)Lox at 2.5 × 106 (MOI 5), 2.5 × 107 (MOI 50), or 5 × 107 PFU/mL (MOI 100), and the amount of cell-associated and extracellular virions was titrated on Vero cells. Each data point represents the mean of the three technical replicates of the combined cell-associated and extracellular titers. The error bars represent the standard deviation.

### IFN-induced protein changes in the cytosol of macrophages

To identify cytosolic macrophage proteins that might foster or restrict HSV-1 capsid functions, we prepared extracts from Mφ_R_ or IFN-induced Mφ_IFN_ to reconstitute capsid-host protein complexes as they might assemble in macrophages (Fig. S1). Using subcellular fractionation and subsequent dialysis (Fig. S2A), we depleted the extracts of nuclei and mitochondria (Fig. S2B; pellet P1), cytoplasmic membranes such as Golgi apparatus, endoplasmic reticulum and plasma membrane (P1, P2), and small metabolites (S2, S3, S4). Furthermore, most of the cytoskeletal tubulin and actin sedimented into the first pellet (P1), while glyceraldehyde 3-phosphate dehydrogenase (GAPDH), a bona-fide cytosolic protein, remained soluble in the supernatants (S1, S2, S2’, S3, S4). Next, we analysed the proteomes of the Mφ_R_ and IFN-induced Mφ_IFN_ cytosols at low ATP/GTP concentration [ATP/GTP^low^] by mass spectrometry (MS; Table S1). We detected 494 (Fig. S2C; black circles) of more than 600 reported IFN-inducible proteins (Rusinova *et al*., 2013). Of those, GALM, COL1A1, LGALS3BP, NT5C3A, IFI44, IFIT2, IFIT3, GBP4, SRP9, IFIT5, DSP, and L3HYPDH were enriched by at least 2-fold in the Mφ_IFN_ cytosol (Fig. S2C; red). These changes might reflect IFN-induced transcriptional or translational regulation, post-translational modification, subcellular localization, or susceptibility to proteolysis, and show that the IFN induction had changed the cytosol proteome of the Mφ_IFN_.

### HSV-1 capsids interact with specific cytosolic macrophage proteins

To search for cytosolic Mφ proteins whose interactions with HSV-1 capsids depend on their surface composition, we generated tegumented viral V_0.1_, V_0.5_, and V_1_ capsids as well as D capsids with a reduced tegumentation (Fig. S1). For this, we lysed extracellular particles released from HSV-1 infected cells with non-ionic detergent to solubilize the envelope proteins and lipids, and in the presence of 0.1, 0.5, or 1 M KCl to modify intra-tegument protein-protein interactions (Anderson *et al*., 2014; Ojala *et al*., 2000; Radtke *et al*., 2010; Radtke *et al*., 2014; Wolfstein *et al*., 2006; Zhang & McKnight, 1993). Furthermore, we dissociated tegument from V_0.1_ capsids by a limited trypsin digestion to generate so-called D capsids. We then incubated similar amounts of different capsid types as calibrated by immunoblot for the major capsid protein VP5 (Fig. S2D) with cytosol at ATP/GTP^low^ from Mφ_R_ or IFN-induced Mφ_IFN_ for 1 h at 37°C. The capsid-host protein complexes assembled *in vitro* were harvested by sedimentation, and their interactomes were determined by quantitative MS (c.f. Fig. S1). As before (Radtke *et al*., 2010; Snijder *et al*., 2017), the protein intensities were normalized across samples to the abundance of the major capsid protein VP5 (Table S2, host; Table S3, viral).

Of 2983 proteins identified (Table S2), we detected 1816 in at least 3 of the 4 replicates in any of the 8 different capsid-host protein complexes. Of those, 598 host proteins bound differentially to one capsid type over another (Table S2; fold change ≥ 2.8; permutation-based FDR ≤ 0.05). The HSV-1 capsids had recruited specifically 279 proteins of Mφ_R_ and 390 of Mφ_IFN_ cytosol of which 71 were shared. Hierarchical clustering analyses of the associated Mφ_R_ or Mφ_IFN_ proteins identified 4 major classes; e.g. one enriched on V over D capsids (Fig. S3A and S3B, top green) and one enriched on D over V capsids (Fig. S3A and S3B, bottom violet). Therefore, we compared the capsid-host interactions of D capsids directly to V_0.1_ (Fig. 2A, 2D), V_0.5_ (Fig. 2B, 2E), or V_1_ (Fig. 2C, 2F) capsids, and identified 82 proteins of Mφ_R_ (Fig. 2A, 2B, 2C) and 141 of Mφ_IFN_ (Fig. 2D, 2E, 2F) with 35 being shared (Table S2; difference ≥ 2.83-fold; FDR ≤ 0.01). The Mφ_R_ capsid-host complexes included 12 and the ones of Mφ_IFN_ 19 proteins listed in the interferome database (Rusinova *et al*., 2013; red in Fig. 2). Gene ontology and pathway enrichment analyses showed that the identified 82 Mφ_R_ (Fig S4) and 141 Mφ_IFN_ (Fig. 3) proteins included many players of innate immunity, intracellular transport, nucleotide and protein metabolism, as well as intracellular signalling. Overall, the host proteomes of V_0.1_ (red) and D (grey) capsids were rather distinct, but more similar for V_0.5_ (blue) and V_1_ (green) capsids (Fig. S4 and Fig. 3). For example, V_0.1_ capsids had recruited specifically the innate immunity proteins PIGR, IGHA1, BPIFA1 and DEFA3, but D capsids LRRFIP1, UFC, C3 and DCD from Mφ_R_ cytosol. In Mφ_IFN_, the D capsids were enriched for C3, C6, IGBP1, UBA5, UBXN1, UBE3A, and RNF123. These data suggest that protein domains displayed on different capsids interacted with specific cytosolic Mφ_R_ or Mφ_IFN_ proteins.

**Figure 2:**
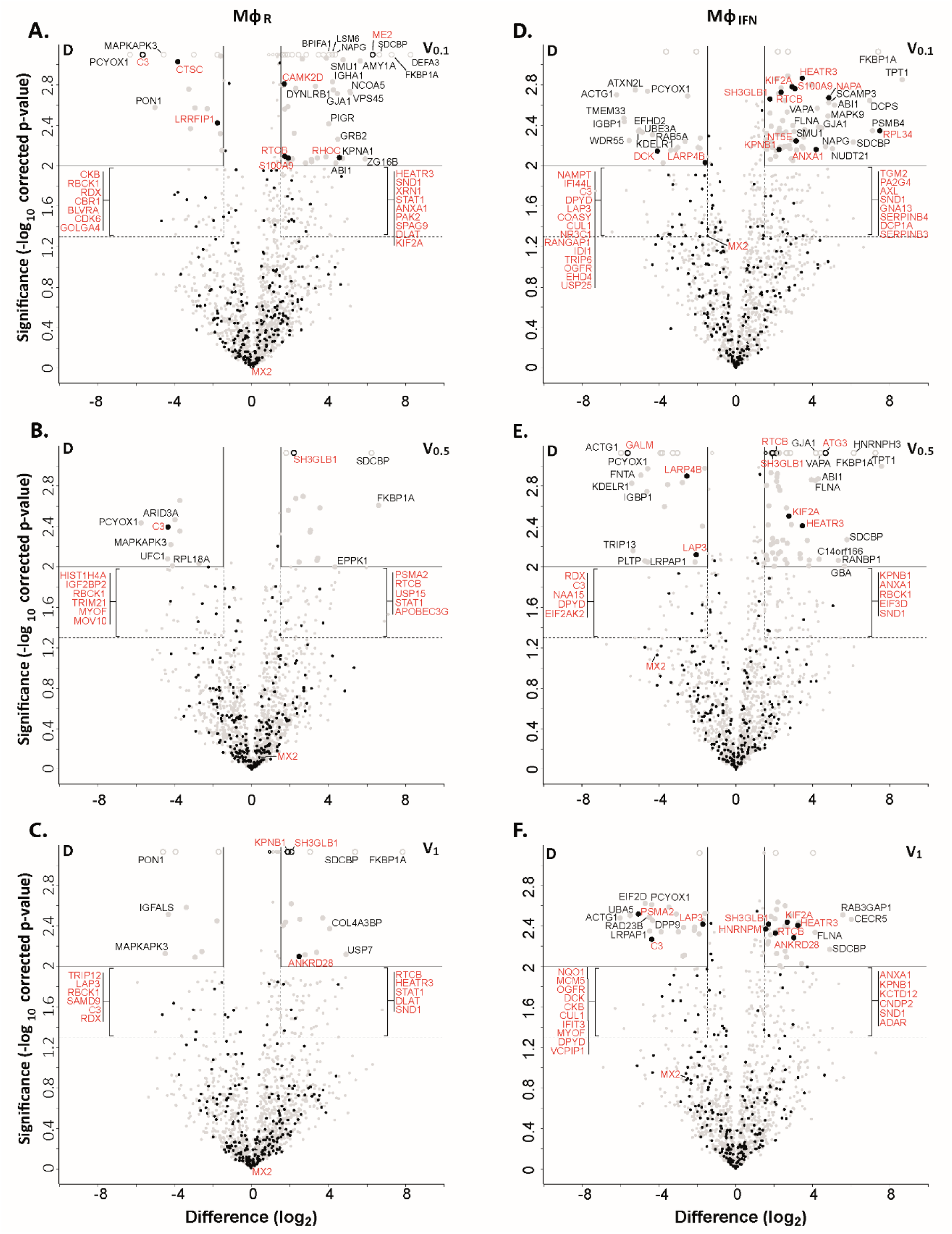
Cytosolic IFN-induced macrophage proteins binding to HSV-1 capsids. Volcano plots of iBAQs counts of proteins identified in capsid-host protein complexes assembled in cytosol from resting THP-1 φ cells (A - C) or treated with interferon-α (D - F) using V_0.1_ (A, D), V_0.5_ (B, E), or V_1_ (C, F) capsids in comparison to D capsids. Proteins identified as highly specific interactions are indicated with larger symbols (log_2_ difference: 1.5; Welch’s t-test, two-tailed, permutation-based FDR ≤ 0.01); those with a log_2_ difference ≥ 4 are annotated. ISGs (interferome.org) are indicated by filled black circles, and are annotated in red if significantly enriched (permutation-based FDR ≤ 0.05, and log_2_ difference ≥ 1.5). Proteins with a q-value = 0 were imputed to - log_10_ q-value = 3.1 (maximum of the graph), and were indicated with empty circles.

**Figure 3:**
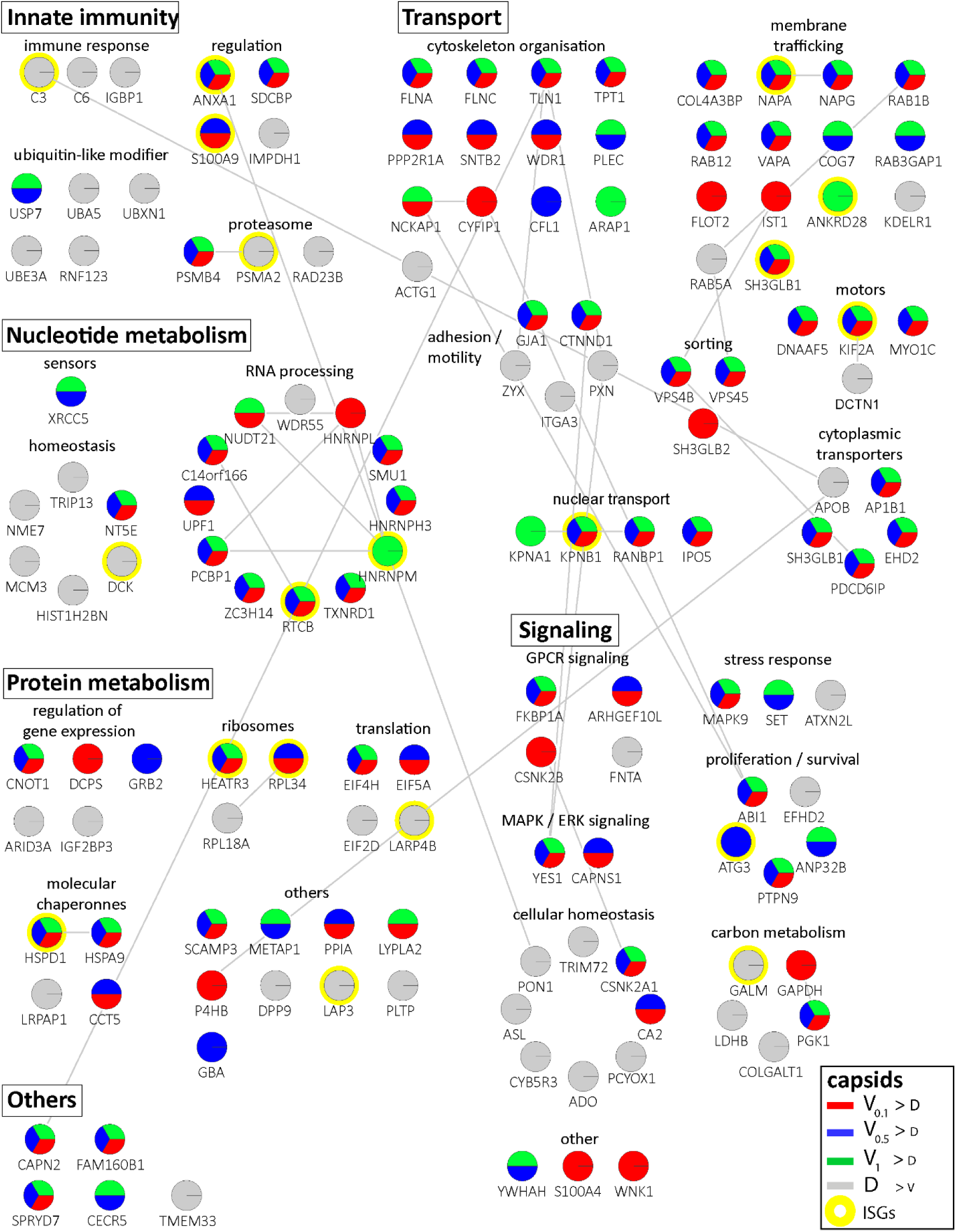
Cytosolic proteins of IFN-induced macrophages binding to HSV-1 capsids. Host proteins from cytosol of IFN-stimulated Mφ_IFN_ (c.f. Fig. 3D, 3E, 3F; abundance log_2_ difference larger than 1.5; significance permutation-based FDR smaller than 0.05) interacting with V_0.1_, V_0.5_, V_1_, or D capsids were assembled into a functional interaction network of known protein-protein-interactions (grey lines; STRING database, confidence score of 0.7), and grouped according to their known functions (Gene Ontology, Pathway analysis). The Pie chart for each protein indicates its relative enrichment on V_0.1_ (red), V_0.5_ (blue), V_1_ (green), or D capsids (grey).

In these assays, the capsids interacted with several proteins already validated to promote or restrict HSV or VZV infection. Examples are the ESCRT-III co-factor VPS4 (Cabrera *et al*., 2019; Crump *et al*., 2007), EIF4H (Page & Read, 2010), the Kif2a subunit of kinesin-13 (Turan *et al*., 2019), the POLR1C subunit of RNA polymerase III (Carter-Timofte *et al*., 2018), the DNA protein kinase PRKDC (Justice *et al*., 2021), and DDX1 (Zhang *et al*., 2011). Moreover, the deubiquitinase USP7 (Rodriguez *et al*., 2020) and the ubiquitin ligases RNF123, TRIM72, UFC1 and UBE3A as well as the proteasome might regulate capsid functionality (Huffmaster *et al*., 2015; Schneider *et al*., 2021) or their degradation(Horan *et al*., 2013; Sun *et al*., 2019). These data show that HSV-1 capsids exposing a different tegument composition recruited specific cytosolic proteins from resting or IFN-induced macrophages.

### HSV-1 capsids recruit specific proteins responding to or regulating type I IFN

We next analysed the Mφ_IFN_ samples in detail as IFN induction had prevented HSV-1 infection completely. We generated cluster maps for the 32 capsid-associated proteins belonging to the GO clusters *Response to type I IFN* or *Regulation of type I IFN production* (Table S2). V capsids recruited DHX9, HSPD1 and FLOT1 as well as proteins involved in the DNA damage response like PRKDC/DNA-PK, XRCC5, and XCCR6 from both, Mφ_R_ and Mφ_IFN_ cytosol (Fig. 4). Interestingly, V capsids bound specifically to STAT1 in Mφ_R_, but to ADAR and IFIT2 in Mφ_IFN_ cytosol. D capsids were enriched for IFI16, OAS2, POLR1C, STAT2, and MxB in Mφ_IFN_ but not in Mφ_R_ (Fig. 4, Fig. S5). Particularly, the discovery of MxB in these capsid-host protein complexes was interesting, as MxB but not its homolog MxA restricts infections of the herpesviruses HSV-1, HSV-2, MCMV, KSHV, and MHV-68, but its mode of action has not been elucidated (Crameri *et al*., 2018; Liu *et al*., 2012; Schilling *et al*., 2018; Vasudevan *et al*., 2018). Therefore, we investigated the interaction of human MxB with HSV-1 capsids further.

**Figure 4:**
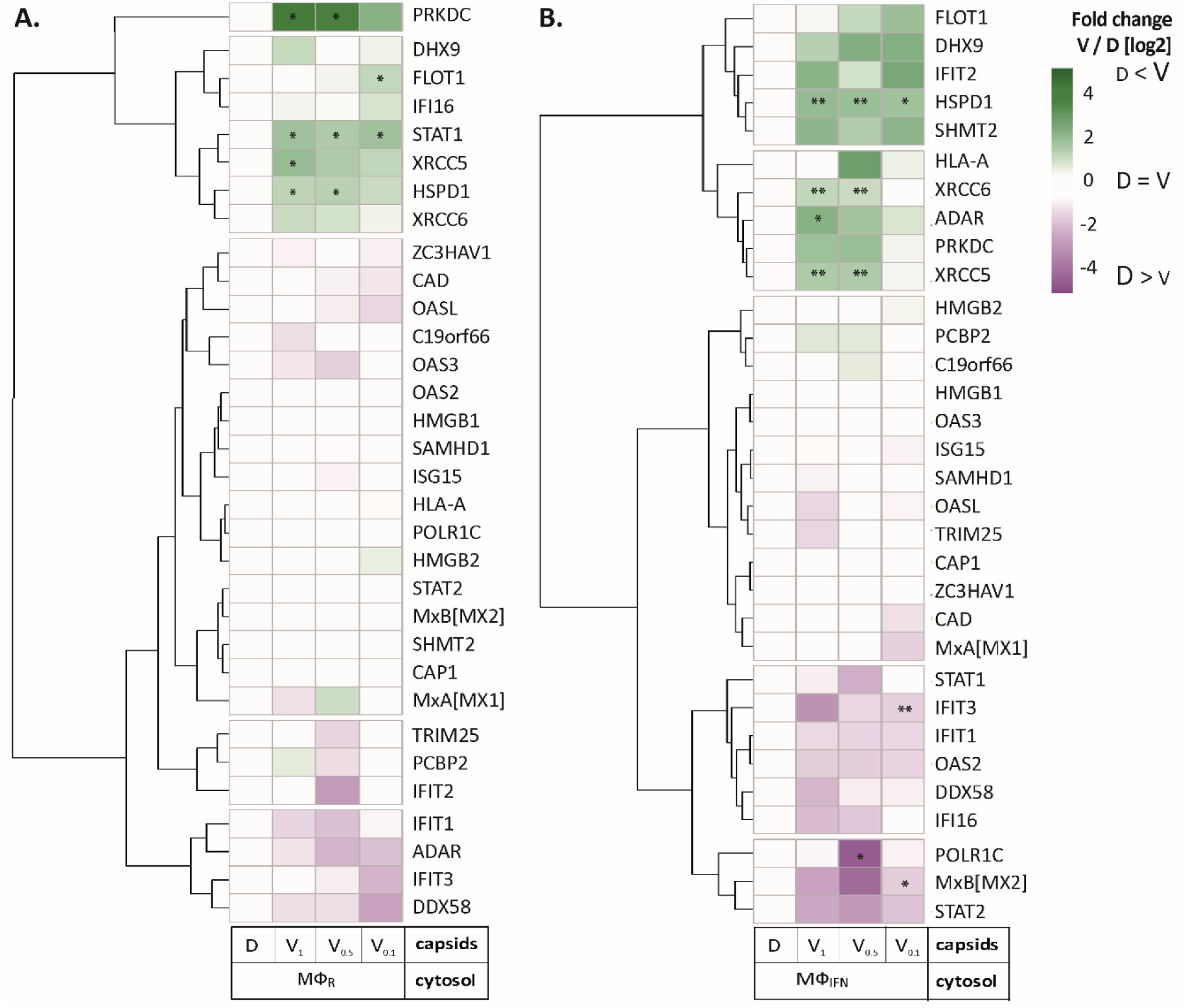
HSV-1 capsids associate with proteins involved in type I IFN response. Unbiased hierarchical clustered heat map showing the log_2_ fold changes of IFN-induced proteins (GO type-I IFN) identified from capsids-host protein sediments from cytosol of resting Mφ, or IFN-induced Mφ_IFN_ macrophages. For each protein, the fold change was calculated based on their abundance (iBAQs) in V_1_, V_0.5_ and V_0.1_ capsids as compared to their abundance in D capsids, using a linear scale from violet being the lowest to dark green being the highest. (*) and (**) design the proteins with an FDR corrected p-value < 0.05 and < 0.01, respectively.

### MxB binds to capsids

We first characterized the MxB fractionation behaviour during the cytosol preparation (Fig. S2). As reported (Goujon *et al*., 2013; Melen *et al*., 1996), MxB was upregulated in IFN-induced Mφ_IFN_. MxB sedimented with nuclei and mitochondria as expected (Cao *et al*., 2020), with cytoplasmic membranes, and possibly filamentous MxB (Alvarez *et al*., 2017) might have been sedimented too. Both, after the addition of ATP and GTP (ATP/GTP^high^) or the hydrolase apyrase (Pilla *et al*., 1996; ATP/GTP^low^), a significant fraction of MxB remained soluble in the cytosol.

Next, we confirmed by immunoblotting that MxB co-sedimented with HSV-1 capsids which had been incubated in cytosols from Mφ_R_ or Mφ_IFN_. In line with the MS results, MxB bound better to D than to V_0.1_, V_0.5_, or V_1_ capsids (Fig. 5A). We next probed authentic nuclear capsids, namely empty A, scaffold-filled B, or DNA-filled C capsids, as well as tegumented V_1_, V_0.5_, V_0.1_ or D capsids with cytosol of A549-MxB(1-715) epithelial cells expressing MxB(1-715). Nuclear A and C as well as V_1_ and D capsids recruited MxB efficiently, while B, V_0.1_ and V_0.5_ capsids bound less MxB (Fig. 5B). MxB did not sediment by itself, and also did not associate with agarose beads used as another sedimentation control (Fig. 5A, 5B). These data indicate that MxB binds to specific structural features on the capsid surface.

**Figure 5:**
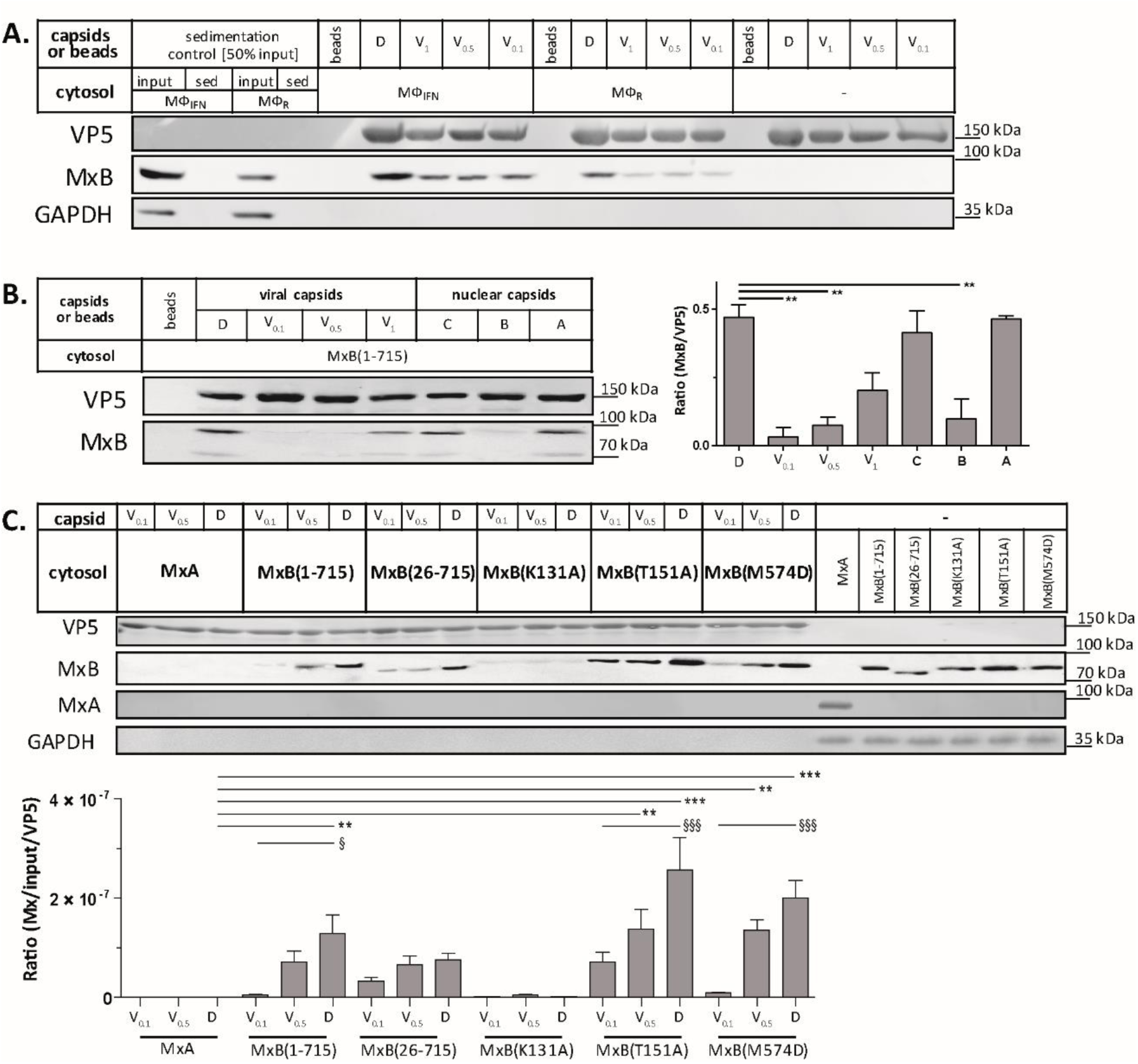
Tegumentation reduces MxB binding to HSV-1 capsids. The binding of MxB to viral V_0.1_, V_0.5_, V_1_ or D, or to nuclear A, B or C capsids was analysed after incubation in 0.2 mg/mL cytosol prepared from (A) THP-1 φ stimulated or not with IFN, or (B-C) A549 cells stably expressing MxA, MxB(1-715) full length, the short MxB(26-715), or MxB mutants defective in GTP-hydrolysis MxB(T151A), GTP-binding and hydrolysis MxB(K131A), or dimerization MxB(M574D). Sedimented capsid-host protein complexes were then analysed by immunoblot for VP5 (capsid), MxB, MxA and GAPDH as a loading control. As control cytosols were sedimented without capsids (A: sed), or with uncoated agarose beads (A, B: beads). The amounts of MxA/MxB found in the capsid-host protein complexes were quantified, and normalized to their respective VP5 levels. Error bars: SEM. summarized from three experiments. One asterisk denotes p<0.05, two asterisks indicate p<0.01 and three asterisks represent p<0.001 as determined by Welch’s t-tests comparisons. **Figure 5A-source data 1-3. Figure 5B-source data 1-3. Figure 5C-source data 1-5.** Full western blot images for the corresponding detail sections shown in Figure 5, as well as raw values for western blot quantifications.

In cells, MxB mediated restriction of herpesvirus replication depends on its N-terminal 25 amino acid residues (NTE), its GTPase activity, and its capacity to form dimers (Crameri *et al*., 2018; Schilling *et al*., 2018; Vasudevan *et al*., 2018). We incubated capsids with cytosols containing Mx<u>A</u>, MxB(1-715), MxB(26-715) (Melen *et al*., 1996; Melen & Julkunen, 1997), MxB(K131A) with reduced GTP binding, MxB(T151A) lacking the GTPase activity, or MxB(M574D) unable to dimerize (Alvarez *et al*., 2017; Fribourgh *et al*., 2014; King *et al*., 2004; Schilling *et al*., 2018). In contrast to MxA, MxB(1-175), MxB(26-715), and MxB(M574D) co-sedimented with capsids to a similar extent. Interestingly, MxB(K131A) did not bind to capsids, while MxB(T151A) bound even stronger (Fig. 5C). These data suggest that conformational changes associated with GTP binding or hydrolysis contribute to MxB interaction with HSV-1 capsids.

### MxB disassembles capsids of alphaherpesviruses

Next, we tested whether MxB might affect HSV-1 capsid stability. While the previous capsid sedimentation assays were performed at ATP/GTP^low^, they suggested that the GTP/GDP state of MxB might modulate its interaction with capsids. To test this experimentally, we supplemented the cytosols with 1 mM GTP, 1 mM ATP, and 7.5 mM creatine phosphate to maintain high ATP/GTP levels [ATP/GTP^high^]. We resuspended sedimented capsid-host protein complexes and applied them onto EM grids (Fig. S1), or we added isolated capsids directly onto EM grids and then placed them on a drop of cytosol to allow the formation of capsid-host protein complexes (Fig. 6A). This direct *on-grid assay* required 50 times fewer capsids than the *sedimentation-resuspension assay* and allowed for time-course analyses. For both, we negatively contrasted the samples with uranyl acetate and analysed them by electron microscopy.

**Figure 6:**
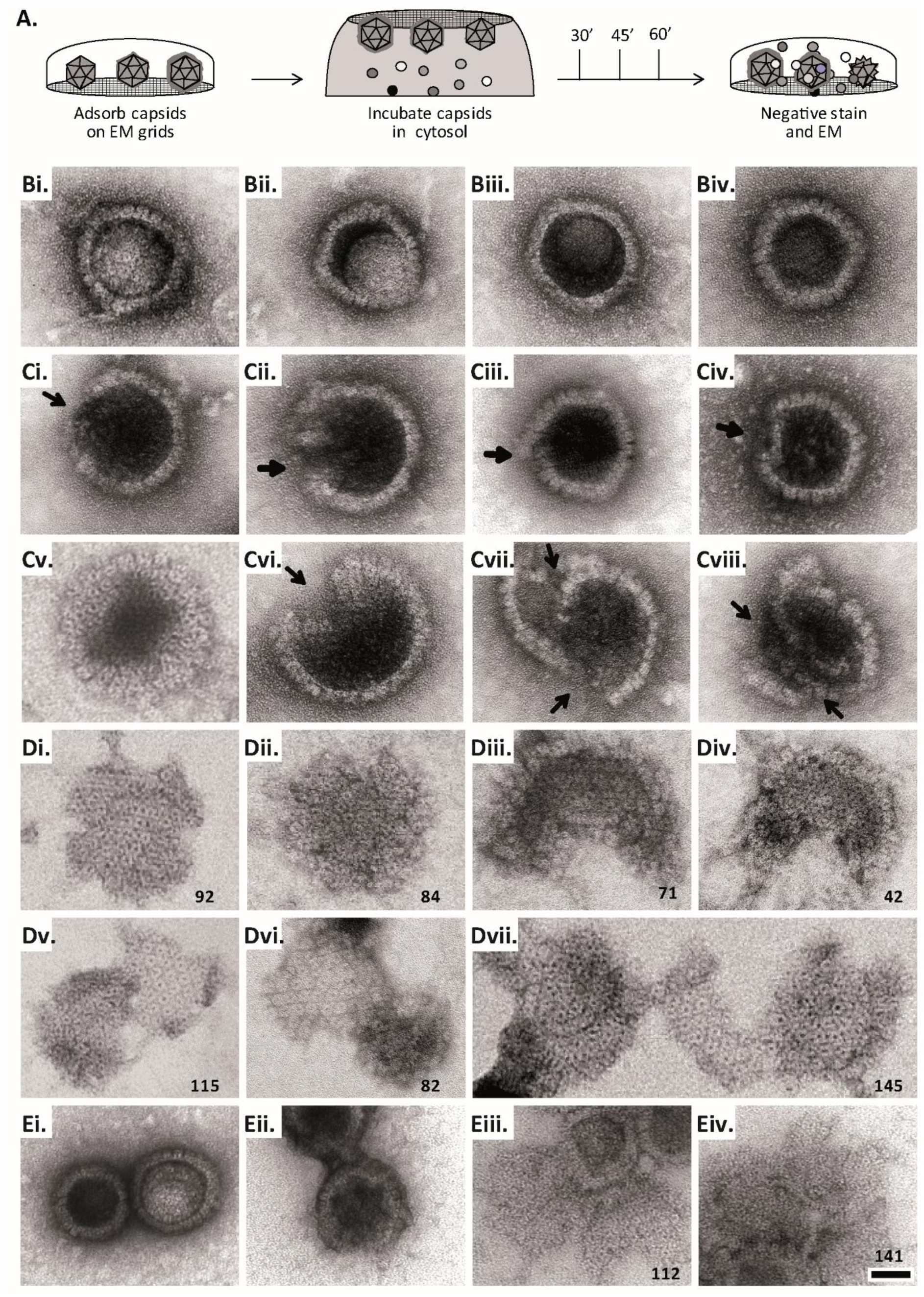
MxB induces disassembly of herpesviral capsids. (A) Experimental design: Capsids were adsorbed onto hydrophilic enhanced carbon-coated EM grids for 20 min at RT. The capsids were incubated in cytosol with ATP/GTP^high^, and the incubation was stopped at different times by extensive washing. The samples were analysed by EM after negative staining with uranyl acetate. (B-D) Capsids after incubation in cytosol derived from rested Mφ or IFN-induced Mφ_IFN_ macrophages, or control or MxB(1-715) A549 expressing cells for 1 h at 37°C, and classified as (B) intact, (C) punched or (D) disassembled flat phenotypes. The number of capsomers per flat particle was counted, and is displayed at the bottom of each figures. (E) Nuclear VZV capsids remain intact (Ei) after incubation in the cytosol of A549 control cells, or but appear punched (Eii) or as flat shells (Eiii, Eiv) after incubation in the cytosol of A549 cells expressing MxB. Scale bar: 50 nm.

When capsids were incubated with cytosol from A549 control cells not containing MxB, we saw mostly intact capsids with an appropriate diameter of about 125 nm, and an intact icosahedral morphology characterized by pentons at the vertices and hexons on the triangular capsid faces (Fig. 6B). The capsids contained genomic DNA as the uranyl acetate used for negative contrast staining had not or only partially entered the capsid lumen. But a treatment with cytosol from IFN-induced Mφ_IFN_ or A549-MxB(1-715) cells dramatically impaired the capsid shell. Based on different MxB induced morphological changes, we classified the capsid structures that we had identified by immunolabeling for capsid proteins (Fig. S6) into three categories. *Intact capsids* (Fig. 6B, Fig. S6A) have an icosahedral morphology and include empty A, scaffold-filled B, and DNA-filled C capsids. *Punched capsids* are characterized by indentations on one or more vertices and an impaired icosahedral shape (Fig. 6C, Fig. S6B). *Flat shells* have completely lost their icosahedral shape (Fig. 6D, Fig. S6C). We estimated the number of capsomers on *flat shells* based on their area, and scored a structure with <100 capsomers as a half capsid and with ≥100 as one capsid (numbers in Fig. 6D). Cytosols containing MxB(1-715) also disassembled capsids of HSV-2 (not shown) or VZV (Fig. 6E) to *punched capsids* and *flat shells*. As MxB induced capsid disassembly of HSV-1, HSV-2 and VZV, these experiments suggest that MxB restricts the infection of herpesviruses by targeting their capsids.

### MxB requires GTP hydrolysis and dimerization to attack herpesviral capsids

Next, we further characterized the capsid disassembly activity of MxB by quantitative electron microscopy. Cytosol from IFN-induced Mφ_IFN_ disassembled more than 80% of the capsids within 1 h while resting Mφ_R_ disassembled only about 40% (Fig. 7A). Cytosol derived from A549 control cells had a minor effect on capsids, while cytosol from A549-MxB(1-715) cells disassembled capsids almost as efficiently as cytosol from Mφ_IFN_. Spiking cytosol from A549 control cells with an increasing percentage of A549-MxB(1-715) cytosol led to an increasing capsid disassembly with a majority of *punched capsids*, at 50% or 66% MxB cytosol, while incubation in pure A549-MxB(1-715) cytosol lead to more than 95% disassembly to mostly *flat shells* within 1 h of incubation (Fig. 7B). We then asked whether MxB had activated other host proteins to mediate capsid disassembly, or whether it was directly responsible. We prepared cytosol from A549-MxB(1-715)-MxB(26-715) expressing both untagged MxB proteins, or from A549-MxB-FLAG expressing MxB(1-715)-FLAG and MxB(26-715)-FLAG. Both cytosols promoted capsid disassembly (MxB; MxB-FLAG in Fig. 7C), but an immunodepletion with anti-FLAG antibodies removed the FLAG-tagged MxB proteins (Fig. S7), and accordingly the disassembly activity from the A549-MxB-FLAG cytosol (MxB-FLAG FT), while the anti-FLAG did neither deplete untagged MxB proteins, nor affect the capsid disassembly activity of the A549-MxB(1-715)-MxB(26-715) cytosol (MxB FT).

**Figure 7:**
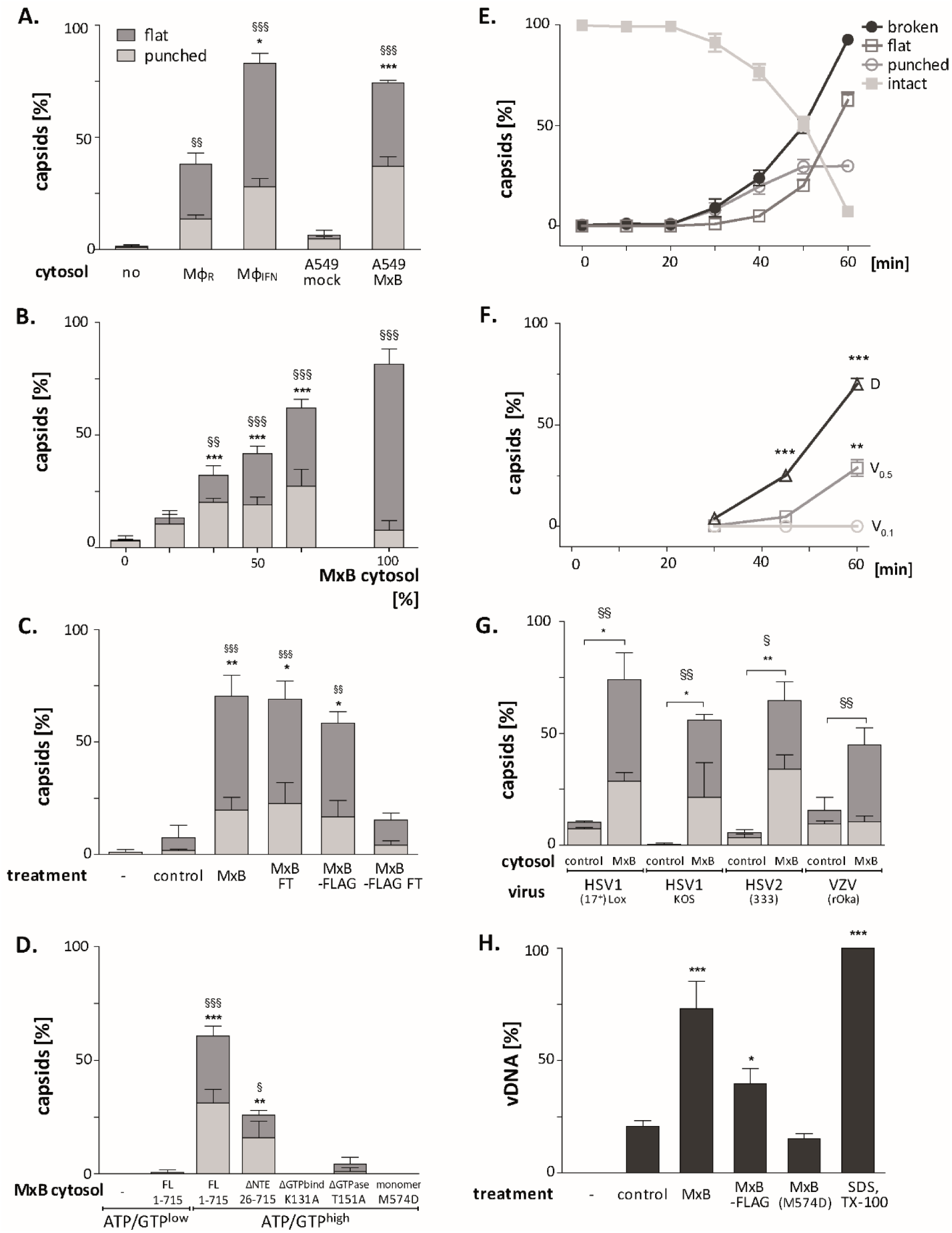
MxB GTP hydrolysis and dimerization required for capsid disassembly and vDNA release of viral genomes. HSV-1 (A-H), HSV-2 (G) or VZV capsids (G) were incubated with cytosol at ATP/GTP^high^ for 1 h or the indicated time (E,F) at 37°C, and classified into *intact, punched* and *flat* capsids by electron microscopy (A-G), or the amount of released viral DNA was measured by qPCR (H). (A) Quantification of *punched* and *flat* D capsid shells in cytosol prepared from rested Mφ or IFN- induced Mφ_IFN_ macrophages, or from control A549 (mock) or A549-MxB(1-715) cells. (B) Increasing amounts of MxB(1-715) [%] were added to control A549 cytosol, and the amount of *punched* and *flat* capsids were quantified after incubation in these mixtures. (C) Cytosols of A549 cells expressing MxB(1-715) and Mx(25-715) or MxB(1-715)-FLAG and MxB(26-715)-FLAG were incubated with anti- FLAG antibodies coupled to magnetic beads, the flow-through fractions (FT) were harvested, capsids were treated with anti-FLAG treated or control cytosols, and the amount of punched and flat capsids were quantified. (D) Capsids were incubated in cytosols prepared from A549 cells expressing full-length (FL) MxB(1-715), MxB(26-715), MxB(K131A), MxB(T151A), or MxB(M574D) at ATP/GTP^low^ or ATP/GTP^high^ levels. (E) Time-course of MxB-induced disassembly of capsids pre-adsorbed onto EM grids, incubated with cytosol from A549-MxB(1-715). (F) Analysis of D, V_0.5_, or V_0.1_ capsids treated with MxB(1-175) cytosol for *broken* (*punched + flat*) capsids after negative stain and EM as described for panel E. (G) Quantification of MxB cytosol disassembly of D capsids of HSV-1(17^+^)Lox, HSV-1(KOS), or HSV-2(333), or nuclear C capsids of VZV, after incubation in cytosol from A549-MxB(1-715) cells. (H) D capsids were incubated with different cytosols for 1 h at 37°C or treated with 1% SDS and 10% Tx-100 only, and the released DNA not protected by capsid shells was quantified by qPCR. Error bars: SEM from 100 capsids in 3 biological replicates. One symbol of *or § denotes p<0.05, two p<0.01, and three p<0.001 as determined in One-way analysis of variance with a Bonferroni post-test, and comparing the relative amounts of (*) *punched* and (§) *flat* capsids, or indicating the differences with the mock treated samples (*).

We next tested at ATP/GTP^high^ the effect of various MxB mutants on HSV-1 capsid stability. While full-length MxB(1-715) induced capsid disassembly, the MxB mutants impaired in GTPase activity (T151A), GTP binding (K131A), or dimerization (M574D) as well as cytosol with MxB at ATP/GTP^low^ did not (Fig. 7D). In contrast, the smaller MxB(26-715) protein lacking the NTE retained about 50% of the capsid disassembly activity. Furthermore, studying the stability of capsids pre-adsorbed *on-grid* in a time-course revealed a lag phase of about 30 min until broken capsids appeared with increasing rate (Fig. 7E). The percentage of *punched capsids* reached a plateau at 50 min, while the amount of *flat shells* continued to increase (Fig. 7E). Further experiments showed that MxB attacked D capsids more efficiently than tegumented V_0.5_ capsids, of which about 70% resisted the MxB attack (Fig. 7F). In contrast, the V_0.1_ capsids seemed to be spared from MxB attack, since no broken capsids appeared within an 1 h treatment. Since MxB restricts infection of several herpesviruses (Crameri *et al*., 2018; Liu *et al*., 2012; Schilling *et al*., 2018; Vasudevan *et al*., 2018), we compared the impact of MxB on D capsids from HSV-1(17^+^)Lox, HSV-1(KOS), HSV-2(333), or on nuclear C capsids from VZV(rOka). Capsids of these human alphaherpesviruses were all susceptible to MxB attack (Fig. 7G).

### MxB attack leads to the release of viral genomes from capsids

Next, we determined how well the capsid shells protected the viral genomes against a DNA nuclease digestion. Capsids released three or two times more viral genomes in cytosols from MxB(1-715) or MxB-FLAG than from control or MxB(M574D) cells (Fig. 7H). Together, these data indicate that the MxB GTPase disassembles the capsid shells and induces a release of viral DNA of several herpesviruses. Our experiments suggest that GTP binding and hydrolysis as well as dimerization contribute to MxB-mediated disassembly of alphaherpesvirus capsids. Its slow start with a lag of about 30 min indicates that the capsid attack might require some nucleating or cooperative reaction to assemble active MxB oligomers or an MxB-containing complex onto capsids.

### Tegument proteins protect against MxB attack

As complete tegumentation shielded V_0.1_ capsids against destruction, while MxB bound to surface features exposed on V_0.5_, A, C and D capsids, we compared the proteomes of the V_0.1_, V_0.5_, V_1_, and D capsids. We calibrated the relative abundances of the 58 HSV-1 proteins detected to the normalized amounts of the major capsid protein VP5. The tegument compositions of V_0.1_, V_0.5_, and V_1_ capsids were similar to each other but different from D capsids (Fig. 8). The bona-fide capsid proteins VP21, VP24, VP22a, VP19c, and VP23 varied little among all capsid types. However, D capsids contain a bit less capsid surface proteins; namely VP26, the capsid specific vertex components (CSVC) pUL17 and pUL25, and to some extent the portal pUL6, and less of the major tegument proteins VP22, VP13/14, VP16, VP11/12 as well as other tegument proteins with ICP0, pUL36 and pUL37 being most susceptible to the trypsin treatment. Overall, there were little differences in the relative tegument protein amounts among V_0.5_ and V_1_ capsids. In contrast, V_0.1_ capsids contained more tegument proteins, e.g. VP13/14, pUS3, pUL41, pUL16, pUS11 and pUL40. All capsid preparations contained traces of membrane proteins and nuclear HSV-1 proteins contributing to DNA replication and packaging (Fig. S8). These data further validated that a treatment with 0.5 or 1 M KCl during the detergent lysis of virions destabilized intra-tegument interactions. Furthermore, the limited trypsin digestion had reduced the capsid proteome further and increased the susceptibility to MxB attack.

**Figure 8:**
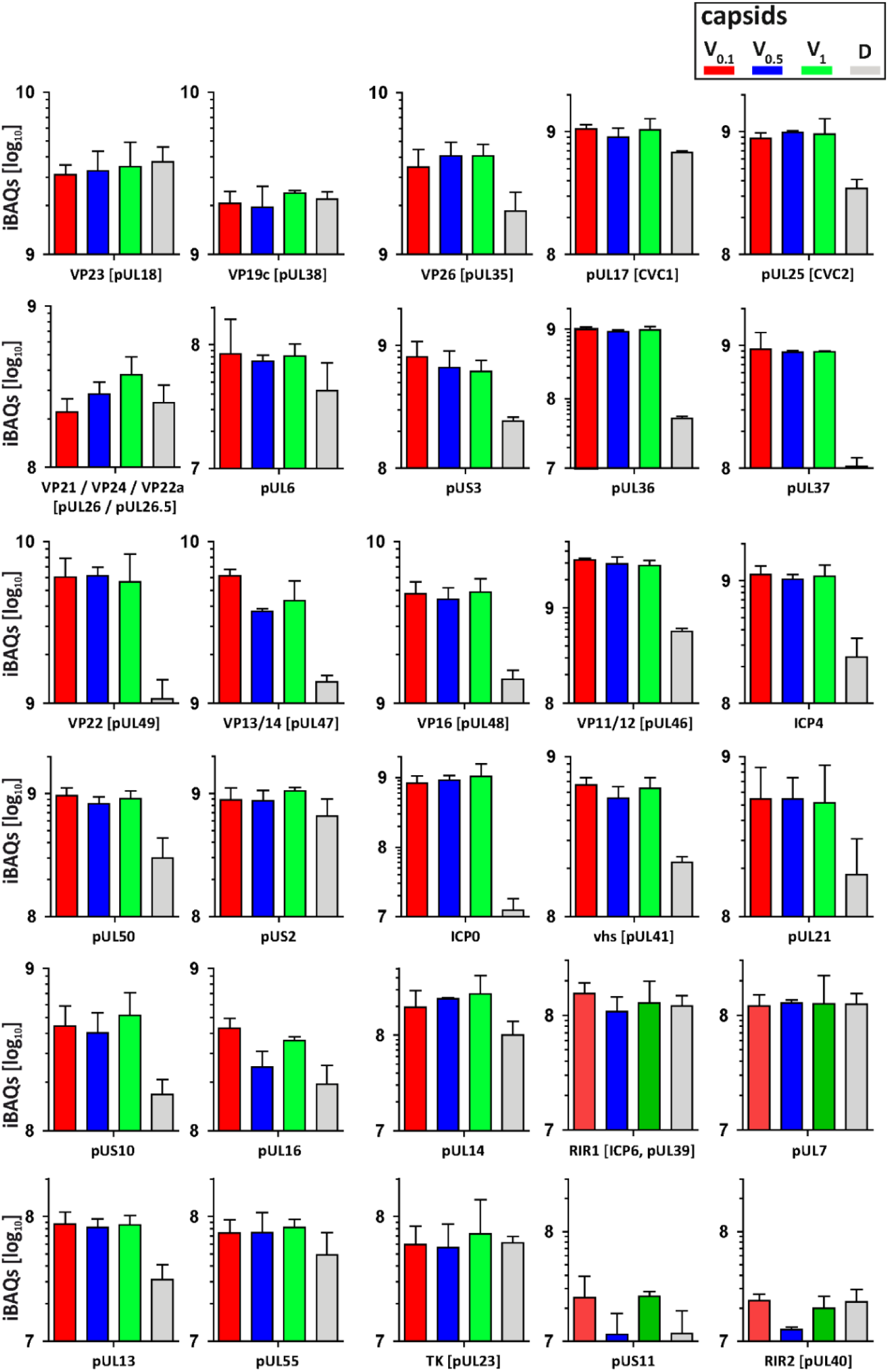
Structural and tegument characterization of V_0.1_, V_0.5_, V_1_, and D capsids. The composition of HSV-1(17^+^)Lox derived V_0.1_ (red), V_0.5_ (blue), V_1_ (green) and D (grey) capsids was analysed by quantitative mass spectrometry in four biological replica. The sum of all the peptides intensities (iBAQ, intensity-based absolute quantification) of each viral protein known to participate in the structure of the capsids was normalized to the one of VP5 and displayed in a bar plot for each viral protein.

## DISCUSSION

Cell-type specific defence mechanisms shape the arms race between proteins restricting or promoting nuclear targeting of incoming viral capsids and viral genome release into the nucleoplasm. We have developed biochemical assays to investigate functional interactions of viral capsids with host cell structures (Radtke *et al*., 2014), and analysed here HSV-1 capsid-host protein complexes assembled in cytosols from resting Mφ_R_ or IFN-induced Mφ_IFN_ cells. We show that the IFN-inducible MxB GTPase bound to alphaherpesviral capsids, most likely to structural features around the capsid vertices, and disassembled herpesvirus capsids in a GTP-dependent fashion, and so that they no longer shielded the viral genomes. Capsid disassembly by MxB could reduce nuclear targeting of incoming capsids and genomes, but stimulate the activation of cytosolic DNA sensors and innate immune responses.

### Cytosolic IFN-induced macrophage proteins binding to HSV-1 capsids

IFN induction prevented HSV-1 infection of Mφ, and increased the cytosolic abundance of at least 12 proteins listed in the interferome database (Rusinova *et al*., 2013). Here, we assembled host protein-capsid complexes from HSV-1 capsids and cytosols of Mφ or Mφ_IFN_ cells as they might also form in cells. While V_0.5_ and V_1_ capsids recruited unique but also common proteins, the proteomes of V_0.1_ and D capsids were more distinct. These results are consistent with immunoelectron microscopy data showing that the surface of distinct V capsid types display different tegument epitopes (Radtke *et al*., 2010), and with cryoelectron tomography data revealing diminishing tegument densities from V_0.1_, V_0.5_, V_1_ capsids to C capsids (Anderson *et al*., 2014). Accordingly, capsids with different tegument composition recruit distinct sets of cytosolic proteins from brain tissue (Radtke *et al*., 2010), or macrophages as shown here. Host proteins may bind to viral proteins in both states, when they are soluble in the cytosol or the nucleoplasm, or when they are associated with capsids. From host proteins shown here to bind to capsids, direct interactions with tegument proteins have already been reported; e.g. USP7 binding to ICP0 (Everett *et al*., 1997) or EIF4H binding to vhs (pUL41; Page & Read, 2010). Furthermore proteins involved in intracellular trafficking or virus assembly associated particularly with tegumented V capsids. For example, importin α5 (*KPNA1*) might mediate capsid targeting to the nuclear pores (Döhner *et al*., 2018; Döhner *et al*., 2021), while RAB1B contributes to the envelopment of cytosolic HSV-1 capsids (Zenner *et al*., 2011).

### MxB binding to alphaherpesviral capsids

In addition to MxB, the host-capsid complexes included other antiviral proteins which in turn might be counteracted by HSV-1 proteins. Several Mφ_IFN_ proteins already know to restrict herpesviruses, e.g. STAT2, POLR1C, IFI16, DDX58 (RIG-I), and OAS2 (Kurt-Jones *et al*., 2017; Lum & Cristea, 2021; Ma *et al*., 2018), bound preferentially to D capsids. As it was not known how MxB might restrict herpesviral infection (Crameri *et al*., 2018; Schilling *et al*., 2018; Vasudevan *et al*., 2018), we investigated its association with capsids further. B capsids are less sturdy and have not undergone the structural changes that stabilize the A and C capsids (Roos *et al*., 2009; Sae-Ueng *et al*., 2014; Snijder *et al*., 2017). Intriguingly, this stabilization depends on the CSVC proteins pUL17 and pUL25 (Sae-Ueng *et al*., 2014; Snijder *et al*., 2017), which are present on B, A, and C capsids (Anderson *et al*., 2014; Radtke *et al*., 2010; Snijder *et al*., 2017). As MxB bound to A, C and D, but not to B capsids, it might recognize surface features formed during capsid stabilization, e.g. matured CSVCs or portals, which are increasingly shielded on tegumented V_1_, V_0.5,_ and V_0.1_ capsids.

Mx<u>A</u> and Mx<u>B</u> GTPases inhibit several viruses by blocking early steps of infection (Haller *et al*., 2015). MxB binding to HIV capsids depends on its N-terminal region (NTR) of about 90 residues and the GTPase domain (Betancor *et al*., 2019; Fricke *et al*., 2014; Smaga *et al*., 2019; Xie *et al*., 2021). Similarly, HSV-1 capsids bound MxB(1-715) and to a lesser extent MxB(26-715). But in contrast to HIV capsids (Betancor *et al*., 2019; Xie *et al*., 2021), HSV-1 capsids recruited also the GTPase deficient MxB(T151A) and the monomeric MxB(M574D). These data indicate that the interaction of MxB with HSV-1 capsids depends on the NTE of 25 residues, its GTP/GDP status, but not on its dimerization.

### MxB induced disassembly of alphaherpesviral capsids

HSV-1 capsid disassembly did not require proteolysis as the cytosols contained protease inhibitors, but may be modulated by other host proteins as there was a considerable lag phase. MxB did not attack fully tegumented V_0.1_ capsids, while V_0.5_ or D capsids were more susceptible. The large tegument protein pUL36 links other tegument proteins to the capsids; it is tightly associated with pUL17 and pUL25 at the CSVCs at the pentons, and it extends towards the 2-fold symmetry axes connecting neighbouring capsid faces (Coller *et al*., 2007; Huet et al. 2016; Liu *et al*., 2019; Newcomb & Brown, 1991; Schipke *et al*., 2012). Our electron microscopy data suggest that MxB attacked the 5-fold symmetry axes as the *punched capsids* had dramatic dents on the capsid vertices. MxB might furthermore attack the portal cap, a cap of HSV1-pUL25 or its homologs in other herpesviruses, which seals the pUL6 portal after DNA packaging is completed (Liu *et al*., 2019; McElwee *et al*., 2018; Döhner et al. 2021; Naniima *et al*., 2021). The high internal capsid pressure due to the negatively charged genome (Bauer *et al*., 2013; Roos *et al*., 2009) could support the MxB attack from the outside. The limited trypsin treatment might have primed the D capsids for disassembly, as they contained less pUL36, pUL17, pUL25, and pUL6 than the V capsids. However, MxB also attacked V_0.5_ capsids that resemble cytosolic capsids during nuclear targeting or after nuclear egress (Ojala *et al*., 2000; Wolfstein *et al*., 2006; Radtke *et al*., 2010; Anderson *et al*., 2014); just not as fast, and not as efficient. Altogether, these results suggest that increasing tegumentation protects incoming and newly assembled capsids, possibly by masking the MxB target structure, or by inhibiting its GTPase cycle.

The MxB-mediated capsid disassembly required its NTE(1-25), GTP hydrolysis, and dimerization. For the homologous MxA GTPase that limits infection of many RNA viruses (Haller *et al*., 2015), Gao *et al*. (2011) proposed a restriction mechanism that involves GTP hydrolysis and a mechano-chemical coupling within ring-like oligomers with the GTPase domains being exposed on their outer diameter (Gao *et al*., 2011). Similarly, MxB can also assemble into helical tubes with the NTE and the GTPase domain oriented outwards (Alvarez *et al*., 2017). Accordingly, MxB monomers and dimers might associate with the capsid vertices and insert between the hexons of neighbouring capsid faces. A further oligomerization of MxB and/or conformational changes associated with GTP hydrolysis might then exert destabilizing forces onto the capsid shells, and ultimately push the capsid faces apart.

### Does MxB induce capsid disassembly in cells?

Future studies need to investigate whether MxB also induces the disassembly of herpesviral capsids in cells. Upon docking of an incoming capsid to a NPC, the pUL25 portal cap is supposed to be displaced, the pUL6 portal to be opened, and the DNA to be ejected from the capsid into the nucleoplasm due to this intramolecular repulsion (Brandariz-Nunez *et al*., 2019; Döhner *et al*., 2021; Ojala *et al*., 2000; Rode *et al*., 2011). In uninfected cells, there is a low amount of constitutively expressed MxB localized at the NPCs (Crameri *et al*., 2018; Kane *et al*., 2018; Melen & Julkunen, 1997), which might dislodge the portal cap and open the capsid portal on the incoming capsid to release the incoming genome into the nucleoplasm.

Crameri *et al*. (2018) proposed that the higher amounts of IFN-induced MxB may block cytosolic capsid transport, genome uncoating at the NPCs, and/or the release of viral genomes into the nucleoplasm, which is consistent with our biochemical data demonstrating MxB binding to HSV-1 capsids. MxB-mediated disassembly of capsids that we report here would further reduce capsid targeting to the NPCs and genome release into the nucleoplasm. Accordingly, there are fewer HSV-1 capsid puncta in MxB expressing cells (Crameri *et al*., 2018). Consistent with our data on capsid disassembly with MxB(26-715), MxB(K131A), or MxB(M574D), restricting the infection of HSV-1, MCMV, and MHV68 also requires the NTE, GTP hydrolysis, and dimerization of MxB (Crameri *et al*., 2018; Schilling *et al*., 2018). Restriction of HIV infection depends also on the NTE, to some extent on the MxB GTPase function, and on its dimerization (Buffone *et al*., 2015; Fricke *et al*., 2014; Goujon *et al*., 2014; Schulte *et al*., 2015; Xie *et al*., 2021). It will be interesting to determine whether MxB only competes for important HIV interactions with promoting host factors (reviewed in Temple *et al*., 2020), or whether it also induces HIV capsid disassembly.

Our data together with Schilling *et al*. (2018) and Crameri *et al*. (2018) suggest that the IFN-inducible MxB restricts HSV-1, HSV-2, VZV, and possibly other herpesviruses, by promoting efficient capsid disassembly. We cannot exclude that a surplus of capsid- and NPC-associated MxB imposes further restrictions on intracellular transport and genome release into the nucleoplasm. However, if MxB(1-715) would disassemble viral capsids before they are oriented properly with their portal towards the NPCs, their genomes would end up in the cytosol and would not be delivered into the nucleoplasm. There are fewer incoming cytoplasmic capsids in cells expressing MxB (Crameri *et al*., 2018), and incoming VP5 is ubiquitinated and degraded by proteasomes in macrophages (Horan *et al*., 2013; Sun *et al*., 2019). Therefore, capsid disassembly intermediates might be degraded in cells, while we could characterize them in our biochemical cell-free assays in which proteases had been blocked.

The viral genomes exposed after MxB-induced capsid disassembly might be degraded by the DNase TREX1 (Sun *et al*., 2019), or stimulate the DNA sensors AIM2, cGAS, or IFI16, and the induction of antiviral host mechanisms. As an inoculation with destabilized HIV-1 capsids leads to an increased activation of the DNA sensor cGAS (Sumner *et al*., 2020), the IFN-induced increased MxB expression might lead to a similar outcome in cells infected with herpesviruses. Accordingly, MxB may not only restrict herpesviruses by capsid disassembly, but also increase the exposure of viral genomes to cytosolic DNA sensors, which in turn would induce an IFN response, inflammation as well as innate and adaptive immune responses. Thus, MxB could be the long sought-after capsid sensor that destroys the sturdy herpesvirus capsids, and possibly HIV cores and other viral capsids, to promote host viral genome sensing.

## MATERIALS AND METHODS

### Cells

All cells were maintained in a humidified incubator at 37°C with 5% CO_2_ and passaged twice per week. BHK-21 (ATCC CCL-10) and Vero cells (ATCC CCL-81) were cultured in MEM Eagle with 1% NEAA (Cytogen, Wetzlar, Germany) and 10% or 7.5% (v/v) FBS, respectively (Good Forte; PAN-Biotech, Aidenbach, Germany). THP-1 cells (ATCC TIB-202; kind gift from Walther Mothes, Yale University, New Haven, USA) were cultured in RPMI Medium 1640 (Thermo Fisher Scientific, Waltham, Massachusetts, United States) with 10% FBS (Thermo Fisher Scientific, Waltham, Massachusetts, United States). THP-1 were stimulated with 100 nM phorbol 12-myristate 13-acetate (PMA; Sigma-Aldrich, Germany) for 48 h and used immediately (Mφ) or after 3 days of rest (Mφ_R_). The cells were cultured with 1000 U/mL human type I IFN-α2a (Mφ_IFN_; R&D Systems, Minneapolis, Minnesota, USA) or left untreated for 16 h.

A549 cells were cultured in DMEM with 10% FCS. In addition to A549 control cells, we used A549 cell lines stably expressing MxB(1-715), MxB(1-715/K131A), MxB(1-715/T151A), MxB(1-715/M574D), MxB(26-715), or MxA(1-662) upon transduction with the respective pLVX vectors with an engineered Kozak sequence to favor expression of the MxB(1-715) over the MxB(26-715) proteins (Schilling et al. 2018). Furthermore, we generated A549-MxBFLAG cells expressing MxB(1-715)FLAG and MxB(26-715)FLAG, both tagged with the FLAG epitope (GACTACAAAGACGATGACGACAAG) at the C-terminus of MxB (GenBAnk NM_002463), and A549-MxB(1-715)-MxB(26-715) cells expressing untagged MxB(1-715) and MxB(26-715) using the pLKOD-Ires-Puro vector (Clontech Takara Bio, Mountain View, United States). MeWo cells (kind gift from Graham Ogg; University of Oxford, Oxford, UK) were cultured in MEM with 10% FCS, NEAA, and 1 mM sodium pyruvate.

### Viruses

Virus stocks of HSV-1(17^+^)Lox (Sandbaumhüter *et al*., 2013), HSV-1 strain KOS (Warner *et al.,* 1998; kind gift from Pat Spear, Northwestern Medical School, Chicago, USA), and HSV-2 strain 333 (Warner *et al.,* 1998; kind gift from Helena Browne, Cambridge University, Cambridge, UK) were prepared as reported before (Döhner *et al*., 2006; Grosche *et al*., 2019). Extracellular particles were harvested from the supernatant of BHK-21 cells infected with 3 to 4 x 10^4^ PFU/mL (MOI of 0.01 PFU/cell) for 2 to 3 days until the cells had detached from the culture flasks, and plaque-titrated on Vero cells. VZV rOka (kind gift from Jeffrey Cohen, NIH, Bethesda, US) was maintained in infected MeWo cells (Cohen & Seidel, 1993; Hertzog *et al*., 2020). After 2 to 4 days, the VZV infected cells as indicated by cytopathic effects were harvested, mixed with naïve MeWo cells at a ratio of 1:4 to 1:8 for continued culture. Aliquots of frozen infected cells were used to inoculate cultures used for capsid preparation.

### HSV-1 infection

THP-1 were seeded at 2.5 x 10^5^ cells per 6-well, treated with 100 nM PMA (Sigma-Aldrich, Germany) for 48 h and used immediately (Mφ) or after 3 days of rest (Mφ_R_). The cells were then induced with 1000 U/mL of IFN-α (Mφ_IFN_) or left untreated for 16 h. On the next day, they were inoculated with HSV-1(17^+^)Lox at 2.5 x 10^6^, 2.5 x 10^7^, or 5 x 10^7^ PFU/mL (MOI of 5, 50, or 100 respectively) in CO_2_-independent medium (Gibco Life Technologies) supplemented with 0.1% (w/v) cell culture grade fatty-acid free bovine serum albumin (BSA; PAA Laboratories GmbH) for 30 min, and then shifted to regular culture medium at 37°C and 5% CO_2._ At the indicated times, the cells and the corresponding media were harvested separately and snap-frozen in liquid nitrogen. These samples as well as and HSV-1 and HSV-2 inocula were titrated on Vero cells (Döhner *et al*., 2006; Grosche *et al*., 2019).

### Preparation of V_0.1_, V_0.5_ and V_1_ and D capsids

Extracellular HSV-1 or HSV-2 particles were harvested by sedimentation at 12,000 rpm for 90 minutes at 4°C (Type 19 rotor, Beckman-Coulter) from the medium of BHK-21 cells (40 x 175 cm² flasks; 2 – 2.5 x 10^7^ cells/flask) infected with 0.01 PFU/cell (2 to 6.7 x 10^4^ PFU/mL) for 2.5 days. The resulting medium pellets (MP) were resuspended in 2 mL of MKT buffer (20 mM MES, 30 mM Tris-HCl, 100 mM KCl, pH 7.4; Grosche et al., 2019; Radtke et al., 2014; Turan et al., 2019), treated with 0.5 mg/mL trypsin (Sigma-Aldrich, Germany) at 37°C for 1 h which was then inactivated with 5 mg/mL trypsin inhibitor from soybean (SBTI; Fluka, Switzerland) for 10 min on ice. These samples were then mixed with an equal volume of 2-fold lysis buffer (2% TX-100, 20 mM MES, 30 mM Tris, pH 7.4, 20 mM DTT, 1x protease inhibitor cocktail [PIs, Roche cOmplete] with 0.2 M, 1 M or 2 M KCl; Radtke et al., 2014). The samples were layered on top of 20% (w/v) sucrose cushions in 20 mM MES, 30 mM Tris, pH 7.4 with 10 mM DTT, PIs with the respective KCl concentration, and sedimented at 110,000 g for 20 min at 4°C (TLA-120.2 rotor, Beckman-Coulter). The supernatants and the cushions containing solubilized viral envelope and tegument proteins were carefully removed. The pellets were resuspended in BRB80 (80 mM PIPES, pH 6.8, 12 mM MgCl_2_, 1 mM EGTA) with 10 mM DTT, PIs, 0.1 U/mL protease-free DNase I (Promega, USA), and 100 mg/mL protease-free RNase (Roth GmbH, Germany) for 1 h at 37°C and then overnight at 4°C. The capsids were sedimented at 110,000 g for 15 min at 4°C (TLA-120.2) and resuspended in capsid binding buffer (CBB: 5% [w/v] sucrose, 20 mM HEPES-KOH, pH 7.3, 80 mM K-acetate, 1 mM EGTA, 2 mM Mg-acetate, 10 mM DTT and PIs) by ultrasound tip sonication at 40 W for about 5 x 5 seconds on ice. Furthermore, we treated V_0.1_ capsids for 40 min at 37°C with 10 µg/mL trypsin in CBB lacking PIs to generate D capsids by limited digestion. After the addition of 5 mg/mL SBTI for 10 min on ice to block the trypsin activity, the D capsids were sedimented at 110,000 x g and 4°C for 15 min (TLA-120.2), and resuspended in CBB with PIs.

### Preparation of nuclear A, B, and C capsids

HSV-1 nuclear capsids were prepared from 40 x 175 cm² flasks with BHK-21 cells infected with 0.01 PFU/cell (3 to 4 x 10^4^ PFU/mL) for about 2.5 days (Anderson *et al*., 2014; Radtke *et al*., 2010; Radtke *et al*., 2014; Snijder *et al*., 2017; Wolfstein *et al*., 2006). VZV nuclear capsids were harvested from infected MeWo cells cultured in 5 to 10 x 175 cm^2^ flasks at maximum syncytia formation but before cell lysis. The cells were harvested, resuspended in MKT buffer (20 mM MES, 30 mM Tris, pH 7.4, 100 mM KCl), snap-frozen, and stored at -80°C. Nuclear A, B, and C capsids were separated by sedimentation at 50,000 x g and 4°C for 80 min (SW40Ti, Beckman Coulter) on linear 20 to 50% sucrose gradients in TKE buffer (20 mM Tris, pH 7.5, 500 mM KCl, 1 mM EDTA; diluted in three volumes of TKE supplemented with 2 mM DTT and PIs (Roche cOmplete). The capsids were sedimented in BSA-coated centrifuge tubes at 110,000 g at 4°C for 20 min (TLA-120.2), resuspended in BRB80 buffer supplemented with 100 mg/mL RNase (Roth, Germany), 0.1 U/mL DNase I (M6101, Promega, USA), 10 mM DTT, and PIs, sedimented again, and resuspended in CBB with PIs.

### Calibration of capsid concentration

To calibrate the amount of capsid equivalents (CAP_eq_) among different experiments, we compared all capsid preparations used in this study with a calibration curve generated from the same starting preparation. The capsids were suspended in sample buffer (1% [w/v] SDS, 50 mM Tris-HCl, pH 6.8, 1% [v/v] β-mercaptoethanol, 5% [v/v] glycerol, PIs [Roche cOmplete]), and adsorbed to nitrocellulose membranes (BioTrace™, Pall Laboratory) using a 48-slot suction device (Bio-DOT-SF, Bio-Rad, Hercules, California, USA). The membranes were probed with a polyclonal rabbit serum raised against purified HSV-1 nuclear capsids (SY4563; Table S4; Döhner *et al.,* 2018) followed by secondary antibodies conjugated to fluorescent infrared dyes (donkey-anti-rabbit IgG-IRDye1 800CW; Table S3), and documented with an Infrared Imaging System (Odyssey, Image Studio Lite Quantification Software, LI-COR Biosciences, Lincoln, Nebraska, USA). MPs harvested from one 175 cm² flasks of BHK-21 cells infected with HSV-1 contained about 0.5 to 1 x 10^9^ PFU/mL, and 0.75 to 1.5 x 10^9^ CAP_eq_/mL. A nuclear HSV-1 capsid fraction prepared from one 175 cm² flask contained about 0.5 to 1 x 10^7^ CAP_eq_ of A capsids, 1 to 2 x 10^7^ CAP_eq_ of B capsids, and 0.5 to 0.75 x 10^7^ CAP_eq_ of C capsids, and a nuclear VZV fraction from one 175 cm² flasks of MeWo cells 2 to 4 x 10^5^ CAP_eq_ of A capsids, 0.5 to 1 x 10^6^ CAP_eq_ of B capsids, and 0.8 to 1.6 x 10^7^ CAP_eq_ of C capsids. Capsid-host protein complexes were assembled *in-solution* using 7.5 x 10^8^ CAP_eq_/condition for MS and immunoblot experiments, and for the *on-grid* electron microscopy assay 2 x 10^7^ CAP_eq_/condition were used.

### Preparation of cytosol

Cytosolic extracts were prepared as described before (Radtke *et al*., 2010; Radtke *et al*., 2014), dialyzed (7K MW cut-off cassettes; Slide-A-Lyzer™, Thermo Scientific), snap-frozen and stored at -80°C. Prior to their use, the cytosols were supplemented with 1 mM ATP, 1 mM GTP, 7 mM creatine phosphate, 5 mM DTT, and PIs (Roche cOmplete), and centrifuged at 130,000 g for 30 min at 4°C (TLA-120.2). We added nocodazole to 25 µM to the cytosols, and left them either untreated (ATP/GTP^high^) or supplemented them with 10 U/mL apyrase (Sigma; ATP/GTP^low^) for 15 min at RT.

### Assembly of capsid-host protein complexes *in-solution*

Capsids were resuspended in CBB and cytosol at a protein concentration of 0.2 mg/mL in an assay volume of 60 µL per sample on a rotating platform at 800 rpm for 1 h at 37°C (c.f. Fig. S1). The capsid-host protein complexes were sedimented through a 30% sucrose cushion at 110,000 g for 20 min at 4°C (TLA-100, Beckman-Coulter), resuspended in CBB by ultrasound tip sonication at 40 W for about 5 x 5 seconds on ice, and analysed by mass spectrometry, immunoblot, or electron microscopy (Radtke *et al*., 2014).

### SDS-PAGE and immunoblot

The samples were lysed in Laemmli buffer (1% [w/v] SDS, 50 mM Tris-HCl, pH 6.8, 1% [v/v] β-mercaptoethanol, 5% [v/v] glycerol, bromophenol blue, PIs [Roche cOmplete]). The proteins were separated on linear 7.5 to 12% or 10 to 15% SDS-PAGE, transferred to methanol-activated PVDF membranes, probed with rabbit or murine primary antibodies (Table S3) and secondary antibodies conjugated to fluorescent infrared dyes (anti-rabbit IgG-IRDye1 800CW; anti-mouse IgG-IRDye1 680RD; Table S3) and documented with an Infrared Imaging System (Odyssey, Image Studio Lite Quantification Software, LI-COR Biosciences, Lincoln, Nebraska, USA).

### Mass spectrometry sample preparation and measurement

Capsid-host protein complexes were analysed by liquid chromatography coupled to tandem mass spectrometry (LC-MS/MS) in four independent biological replicates. The samples were resuspended in hot Laemmli buffer and separated in NuPAGE™ 4 to 12% Bis-Tris protein gels (Invitrogen) before *in-*gel digestion. Briefly, proteins were fixed and stained by Coomassie solution (0.4% G250, 30% methanol, 10% acetic acid). Sample lanes were excised, destained (50% ethanol, 25 mM ammonium bi-carbonate), dehydrated with 100% ethanol and dried using a SpeedVac centrifuge (Eppendorf, Concentrator plus). Gel pieces were rehydrated in trypsin solution (1/50 [w/w] trypsin/protein) overnight at 37°C. Tryptic peptides were extracted in extraction buffer (3% trifluoroacetic acid, 30% acetonitrile), dried using a SpeedVac centrifuge, resuspended in 2 M Tris-HCl buffer before reduction and alkylation using 10 mM Tris(2-carboxyethyl)phosphine, 40 mM 2-Chloroacetamide in 25 mM Tris-HCl pH 8.5. The peptides were purified, concentrated on StageTips with three C18 Empore filter discs (3M), separated on a liquid chromatography instrument, and analysed by mass spectrometry (EASY-nLC 1200 system on an LTQ-Orbitrap XL; Thermo Fisher Scientific) as described before (Hubel *et al*., 2019). Peptides were loaded on a 20 cm reverse-phase analytical column (75 μm column diameter; ReproSil-Pur C18-AQ 1.9 μm resin; Dr. Maisch) and separated using a 120 min acetonitrile gradient. The mass spectrometer was operated in Data-Dependent Analysis mode (DDA, XCalibur software v.3.0, Thermo Fisher).

### Mass-spectrometry data analysis

Raw files were processed with MaxQuant using iBAQ quantification and Match Between Runs option, and the protein groups were filtered with Perseus for reverse identification, modification site only identification, and MaxQuant contaminant list (https://maxquant.net/maxquant/, v1.6.2.10; https://maxquant.net/perseus/, v1.6.5.0; Cox & Mann, 2008; Tyanova *et al*., 2016a; Tyanova *et al*., 2016b). The iBAQ intensities were normalized across all samples to the overall median intensity of the HSV-1 capsid protein VP5. Cytosol and beads incubated with cytosol samples were normalized to all proteins detected in at least three replicates in each condition. Significant differences between given conditions were determined by a two-sided Welch t-test on protein groups present in three replicates of at least one condition, followed by permutation-based FDR statistics (250 permutations), using an absolute log_2_ difference cut-off of 1.5 and an FDR cut-off of 0.01. To characterize the IFN induction, we annotated proteins reported as being induced by IFN type-I as *ISGs* proteins (InterferomeDB, > 2x change; http://www.interferome.org/interferome/home.jspx; Rusinova *et al*., 2013). We used the Fisher’3 exact test against *ISGs* proteins as well as all Gene Ontology (GO) terms for enrichment analysis of proteins upregulated in IFN-induced Mφ_IFN_ cytosol over Mφ_R_ cytosol (log_2_ difference ≥ 1.5; unadjusted p-value < 0.05). The data were summarized in volcano or bar plots (GraphPad Prism v5.0, https://www.graphpad.com/; Perseus v1.6.5.0; Tyanova *et al*., 2016b).

### Interaction Network Assembly

We focused our analysis on proteins that showed specific differences from one capsid preparation to the other, within the same cytosol preparation, and considered host proteins with an enrichment higher than 1.5 log_2_ fold changes and a permutation-based FDR < 0.05 as specifically enriched. To visualize enrichment among different capsid-host protein complexes, we generated integrative networks using Cytoscape (http://www.cytoscape.org/; v3.7.2) and STRING (confidence score: 0.7; Szklarczyk *et al*., 2019). STRING uses a combination of databases on co-expression, conserved occurrences, GO terms and Kyoto Encyclopedia of Genes and Genomes (KEGG; https://www.genome.jp/kegg/; Kanehisa & Goto, 2000; Kanehisa, 2019; Kanehisa *et al*., 2021). To assemble pathway enrichments, we used DAVID, a Database for Annotation, Visualization and Integrated Discovery (https://david.ncifcrf.gov/home.jsp; v6.8; Huang da *et al*., 2009a; Huang da *et al*., 2009b) and the Cytoscape plug-ins ClueGO and CluePedia (http://apps.cytoscape.org/apps/cluego, v2.5.7; http://apps.cytoscape.org/apps/cluepedia, v1.5.7; Bindea *et al*., 2009; Bindea *et al*., 2013).

### Electron microscopy

Capsid-host protein complexes were assembled at ATP/GTP^high^ in solution, harvested by ultracentrifugation, resuspended in CBB, and adsorbed onto enhanced hydrophilicity-400 mesh formvar- and carbon-coated copper grids (Stork Veco, The Netherlands; Radtke *et al*., 2010; Roos *et al*., 2009). Moreover, capsids at a concentration of 1 x 10^7^ CAP_eq_/mL were adsorbed directly for 20 min at RT onto the grids. The grids were incubated on a 10 µL drop of cytosol with a protein concentration of 0.2 mg/mL and ATP/GTP^high^ in a humid chamber for 1 h at 37°C. The samples were left untreated or labelled with anti-VP5 (pAb NC-1) and protein-A gold (10 nm diameter; Cell Microscopy Centre, Utrecht School of Medicine, The Netherlands). For both protocols, the grids were washed with PBS and ddH_2_O, contrasted with 2% uranyl acetate at pH 4.4, air dried, and analysed by transmission electron microscopy (Morgani or Tecnai; FEI, Einthoven, The Netherlands). The capsid morphology was evaluated for about 100 structures/assay from about 15 randomly selected images of 2.7 µm² of three biological replicates. We classified capsomer-containing structures as *punched*, if they lacked one or more of their vertices but still had an icosahedral shape, and as *flat shells*, if they lacked the icosahedral shape but contained capsomers, and scored them as one capsid equivalent structure if they contained more than 100 capsomers.

### Capsid DNA uncoating assay

D capsids were incubated with cytosols from A549-control, A549-MxB(1-715), A549-MxB(M574D), or A549-MxB-FLAG for 1 h at 37°C or treated for 5 min with 1% SDS followed by 10 min with 10% TX-100(Ojala *et al*., 2000). The viral genomes released during the assay were degraded by adding 50 U/mL of benzonase for 1 h at 37°C, and the remaining protected DNA was purified with the DNA Blood Mini Kit (Qiagen, Hilden, Germany) and quantified by real-time PCR on a qTower^3^ (Analytik Jena, Jena, Germany). The SYBR Green assay was performed with the Luna Universal qPCR Master Mix (NEB, Ipswich, MA, USA) according to the manufacturer’s instructions with primers specific for HSV-1 gB (UL27 gene) (HSV1_2 SYBR fwd: 5’- gtagccgtaaaacggggaca-3’ and HSV1_2 SYBR rev: 5’-ccgacctcaagtacaacccc-3’; Engelmann *et al*., 2008). Standards and samples were run in triplicates and results expressed as % released viral DNA with the SDS/Tx-100 treatment normalized to 100%.

### Quantification and statistical analyses

We performed Welch’s t-testing, Kruskal-Wallis H-testing, Friedman and one-way analyses of variance with a Dunns or Bonferroni post-testing (GraphPad Prism v5.0; https://www.graphpad.com/).

### Data availability

The datasets produced in this study are available at PRIDE (PXD028276; http://www.ebi.ac.uk/pride). The published article will include all datasets generated and analysed.

## ACKNOWLEDGMENT

We thank Katinka Döhner and Franziska Hüsers (Institute of Virology, Hannover Medical School) as well as Miriam Schilling (University of Oxford, UK) for many constructive discussions and feedback on the manuscript, and Jasper Götting (Institute of Virology, Hannover Medical School) for support on bioinformatics analyses. We are grateful to Ari Helenius (ETH Zürich, Switzerland), Graham Ogg (University of Oxford, UK), Gary Cohen (University of Pennsylvania, USA),Helena Browne (Cambridge University, UK), Jay Brown (University of Virginia, USA), Jeffrey Cohen (NIH, Bethesda, USA), Pat Spear (Northwestern Medical School, USA), and Roselyn Eisenberg (University of Pennsylvania, USA) for their generous donation of virus strains and invaluable antibodies.

Our research was supported by the EU 7^th^ framework (Marie-Curie Actions ITN-EDGE; https://ec.europa.eu/research/mariecurieactions/about/innovative-training-networks_en, H2020-EU.1.3.1, #675278 to JR, AP, and BS), the UK MRC (core funding of theMedical Research Council Human Immunology Unit to JR), the NIH (NIGMS, GM114141 to IMC), an EU ERC consolidator grant (ERC-CoG ProDAP 817798 to AP), and the German Research Foundation (http://www.dfg.de/; PI1084/3, PI1084/4, PI1084/5, TRR179, and TRR237 to AP; KO1579/13 to GK; CRC900 C2 158989968, EXC62 REBIRTH 24102914, EXC2155 RESIST 390874280, SO403/6 to BS). The funders had no role in study design, data collection and analysis, decision to publish, or preparation of the manuscript.

## AUTHOR CONTRIBUTIONS

MCS and BS conceived and wrote the article. MCS, VG, APir, AHM, TG, FA, IC, APic, and BS contributed to the development of methodology. MCS, VG, SW, and ARN performed experiments and curated the respective data. MCS, VG, APir, APic, and BS analysed the data. MCS, AHM, JH, AB, APo, UP, and SW produced the resources used in this study. FA, SW, TG, RB, IC, JR, and GK contributed to the analysis of the data and the discussion of the content. All authors reviewed and edited the manuscript before submission.

## TABLE LEGENDS

***Supplementary Table S1: Host proteins in THP-1 cytosols.*** Intensity-Based Absolute Quantitation (iBAQ) counts of the host proteins identified in the proteomic analysis of the cytosolic extracts prepared from rested or IFN-induced THP-1 φ cytosol. Statistical analyses were performed with a Welch’s t-test. The following cut-offs were set for differentially-expressed proteins: permutation-based false-discovery rate ≤ 0.05 and |log_2_ fold-change| ≥ 0.5. The protein groups were filtered to keep only the intensities measured in at least three out of four replicates per condition. Gene Ontology knowledge was used to reference the proteins previously described as induced by interferon.

***Supplementary Table S2: Host proteins in capsid-host protein complexes.*** Intensity-Based Absolute Quantitation (iBAQ) counts of host proteins identified in the V_0.1_, V_0.5_, V_1_ and D capsid-host protein complexes assembled in rested or IFN-induced THP-1 φ cytosol. Statistical analyses were performed with a Welch’s t-test. The following cut-offs were set for differentially expressed proteins: permutation-based false-discovery rate ≤ 0.05 and a |log_2_ fold-change ≥ 1.5|. The protein groups were filtered to keep only those with intensities measured in at least three out of four replicates, in at least one condition. “Interaction significance” column indicates the proteins considered as specific interactors.

***Supplementary Table S3: Viral proteins in capsid-host protein complexes.*** Intensity-based absolute quantification (iBAQ) counts of HSV-1(17^+^)Lox viral proteins from isolated V_0.1_, V_0.5_, V_1_ and D capsids (A) normalized to the intensity of the major capsid protein VP5, (B) unnormalized LFQ intensities. The viral proteins were filtered to keep only those with intensities measured in at least three out of four replicates, in at least one condition.

**Supplementary Table S4:** List of Antibodies. mAb: monoclonal antibody. pAb: polyclonal antibody.

| Antigen | Name | species, type | Source and Reference |
| --- | --- | --- | --- |
| **HSV-1 proteins** |  |  |  |
| capsid | SY4563 | rabbit pAb | (Döhner *et al.*, 2018) |
| VP5 | NC-1 | rabbit mAb | Gary Cohen & Roselyn Eisenberg, University of Pennsylvania, Philadelphia, USA; (Cohen, G. H. *et al.*, 1980) |
| **Host proteins** |  |  |  |
| nuclear pore | mAb414 | mouse mAb | ab24609, Abcam |
| p230 | p230 | mouse mAb | 611280, BD Biosciences |
| E-cadherin | α-E-cadherin | mouse mAb | C37020;610404, BD Transduction Laboratories |
| calnexin | α-calnexin | rabbit pAb | Ari Helenius, ETH Zürich, Switzerland; (Hammond & Helenius, 1994) |
| Tom20 | F-10 | mouse mAb | Sc-17764, Santa Cruz Biotechnology |
| GAPDH | 14C10 | rabbit pAb | 2118S, Cell Signaling (NEB) |
| MxA/MxB | M143 | mouse mAb | (Flohr *et al.*, 1999) |
| MxA | α-Mx1 | rabbit pAb | ab207414, Abcam |
| MxB | α-Mx2 | rabbit pAb | NBP1-81018, Novus Biological |
| FLAG | ANTI-FLAG | rabbit pAb | F7425, Sigma-Aldrich |

**Supplementary Figure S1:**
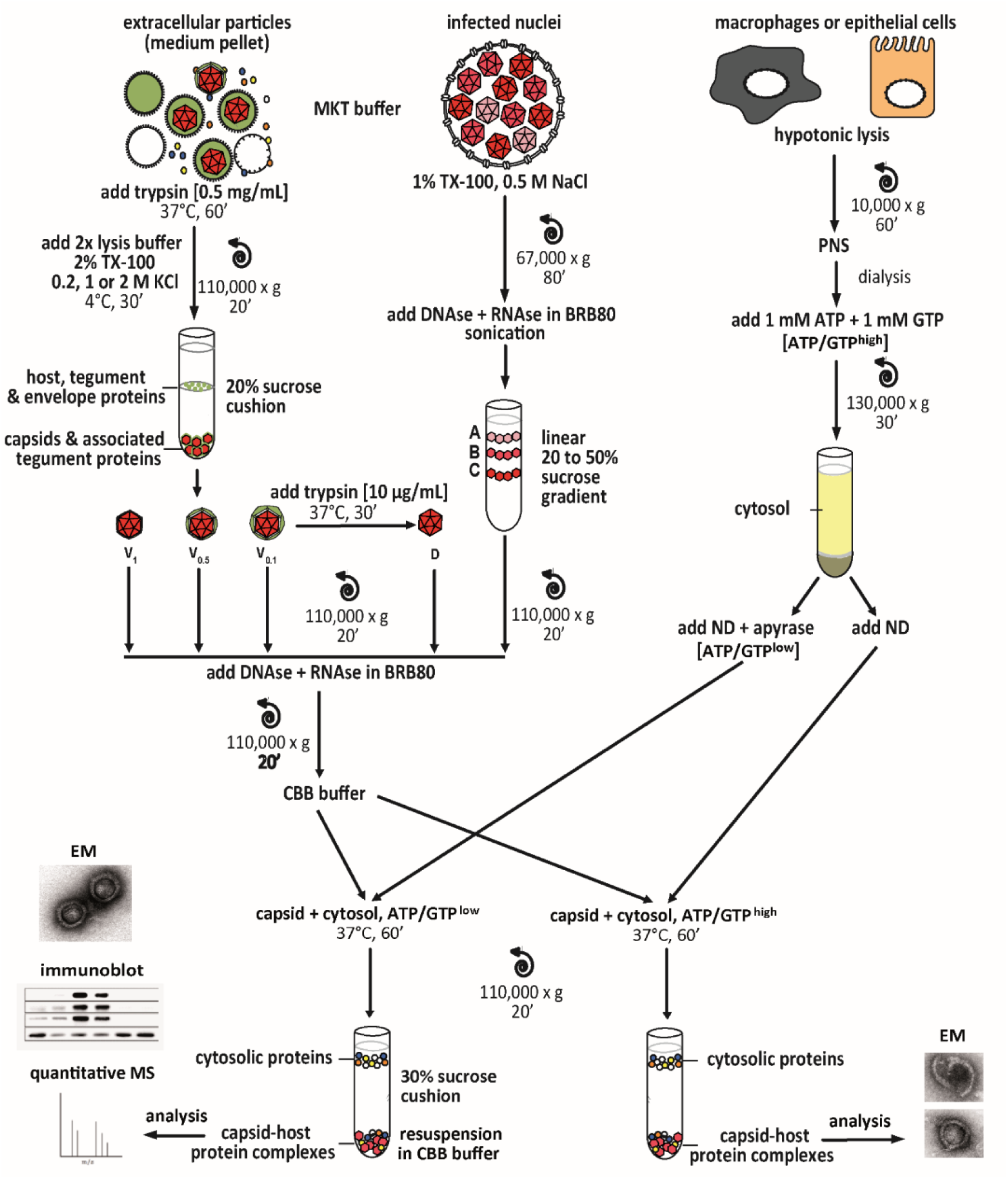
Experimental strategy to generate host protein-capsid complexes. Tegumented viral V_0.1_, V_0.5_, or V_1_ capsids (red) were isolated from extracellular particles released from BHK-21 cells infected with HSV-1(17^+^)Lox. They were lysed in 1% Triton X-100 to solubilize the viral envelope, and to extract different amounts of tegument (green) in the presence of 0.1 M, 0.5 M or 1 M KCl. D capsids were generated from V_0.1_ capsids by mild trypsin digestion. These different capsid types were purified through sucrose cushions. Tegument-free nuclear A, B, and C capsids were isolated from the nuclei of BHK cells infected with HSV-1(17^+^)Lox by gradient sedimentation. The capsids were resuspended in BRB80 buffer, treated with benzonase to degrade DNA and RNA, sedimented again, and incubated with cytosol fractions (yellow) from control or IFN-induced macrophages (THP-1 φ) or epithelial A549 cells. After sedimentation through sucrose cushions, the capsid-host protein complexes were analysed by mass spectrometry (MS), immunoblot, or electron microscopy (EM). PNS, post-nuclear-supernatant; ND, nocodazole.

**Supplementary Figure S2:**
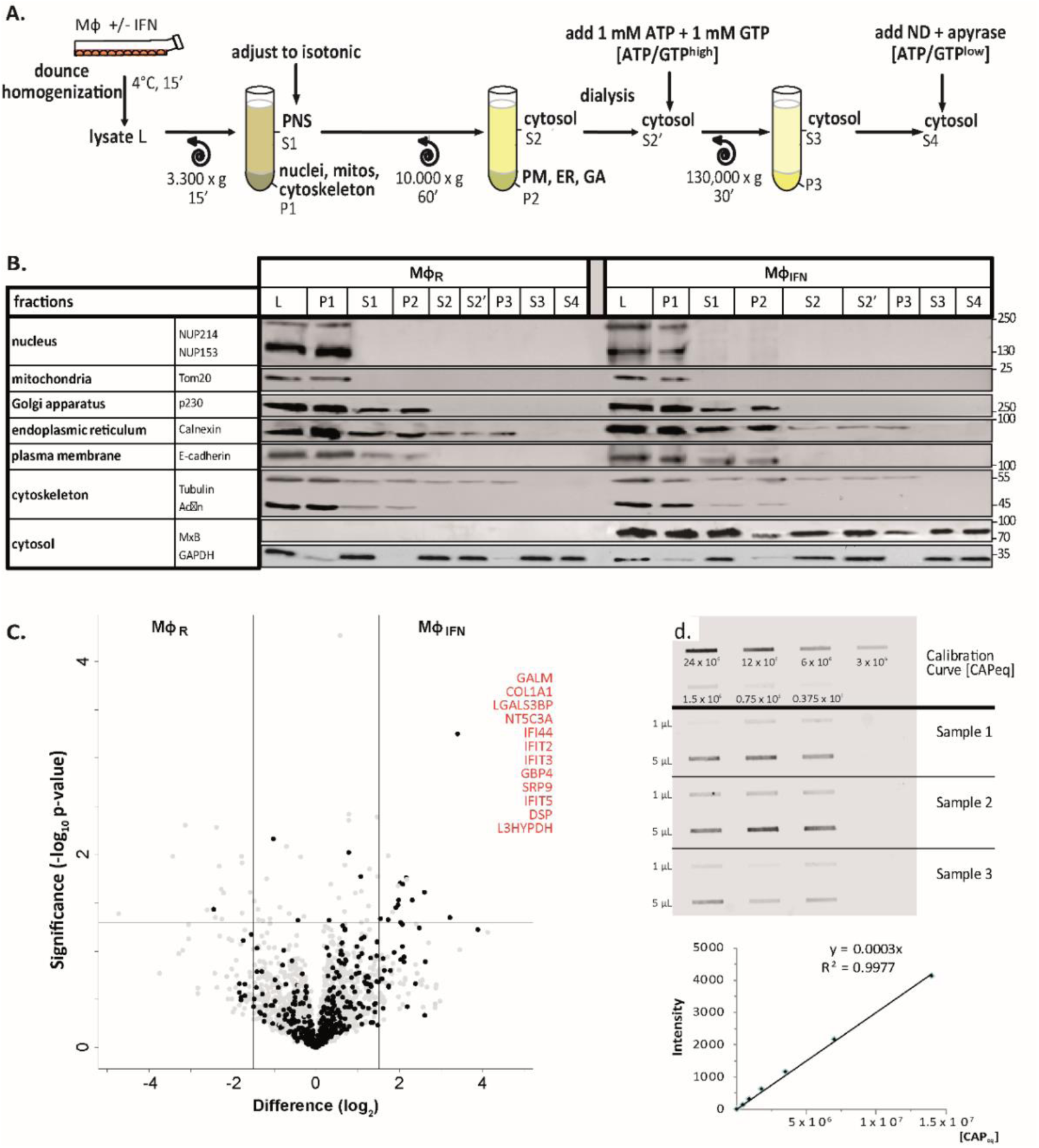
Characterization of cytosolic extracts and calibration of capsids. (A) Cytosols were prepared from rested Mφ_R_ or IFN-induced Mφ_IFN_ macrophage cells. After swelling in hypotonic buffer, the cells were homogenized (L), and nuclei and mitochondria were sedimented (P1). The post-nuclear supernatant (S1) was adjusted to isotonic salt concentration, and centrifuged to sediment membrane compartments (P2), like the PM, ER and GA. To control the nucleotide concentration, the cytosols (S2) were dialyzed against a 7 kDa membrane prior to the addition of an ATP regeneration system (S2’). The remaining actin filaments and microtubules were sedimented in P3 to obtain a soluble cytosol fraction (S3). To reduce ATP and GTP levels, some cytosols were treated with 10 U/mL of apyrase (S4). Nocodazole (ND) was added to prevent polymerization and sedimentation of microtubules. (B) All fractions generated were analysed by immunoblot for the respective compartment marker proteins as indicated. Nup, nucleoporins. (C) Volcano plot summarizing the effect of IFN induction on the cytosol proteome. ISGs associated with the interferomeDB were enriched in cytosol from Mφ_IFN_ as compared to Mφ_R_ with an FDR of 7.96 x 10^-7^ and an FC ≥ 2 in at least 1 experiment (Fisher’s exact test). IFN-inducible proteins are indicated by black circles, and those with an abundance log_2_ difference ≥ 1.5 (vertical lines), and an uncorrected p-value < 0.05 (horizontal line) are labelled in red. (D) The slot blot used for the estimation of capsid concentrations (capsids equivalent; CAP_eq_) of all preparations was labeled with anti-capsid antibodies (rabbit pAb SY4563) and adjusted to a calibration curve of a standard preparation.

**Supplementary Figure S3A:**
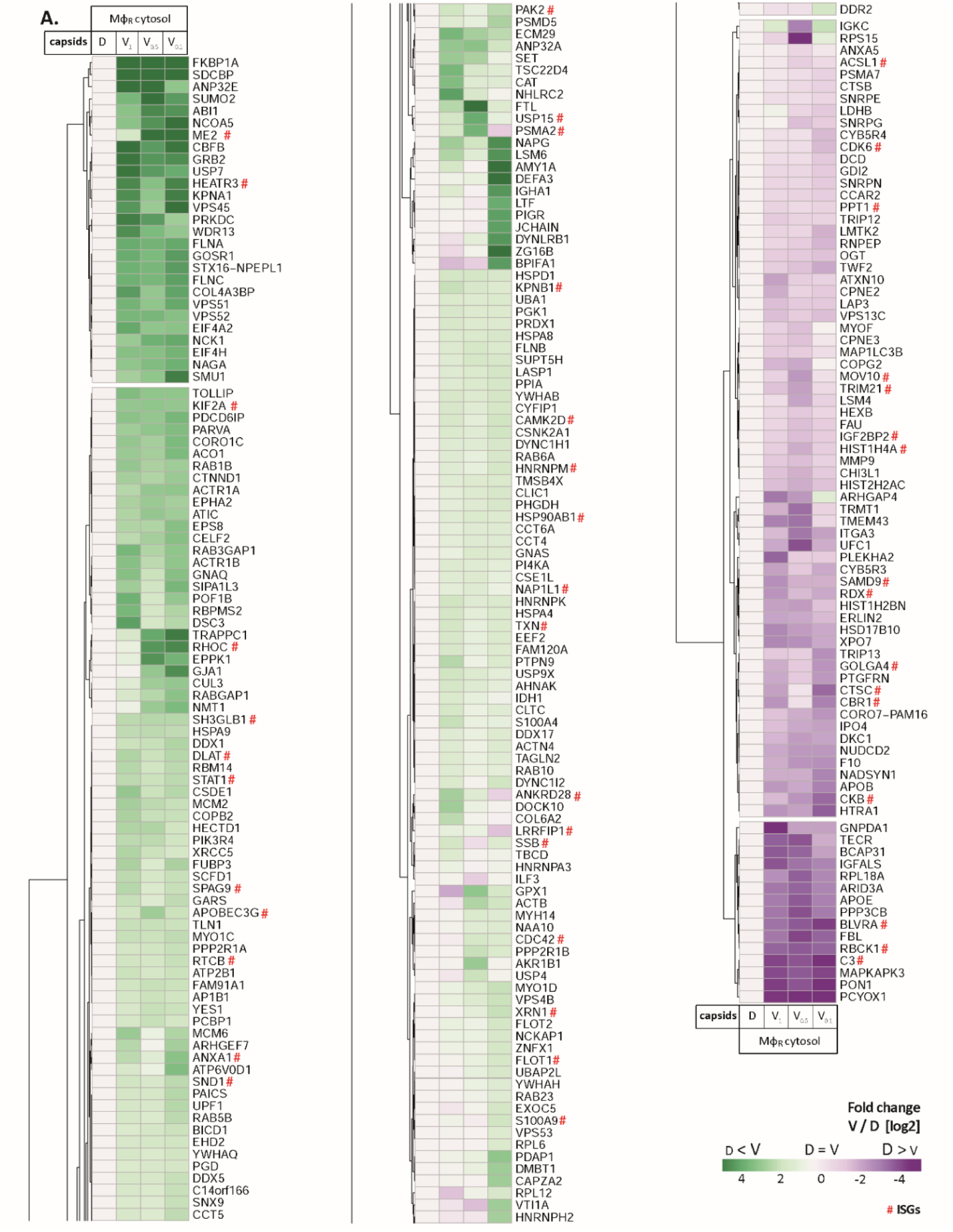
HSV-1 capsids interactomes. Unbiased hierarchical clustered heat map showing the log_2_ fold changes of host proteins identified from capsids-host protein sediments (c.f. Fig. 2; abundance log_2_ difference larger than 1; significance permutation-based FDR smaller than 0.05) from (A) cytosol of resting Mφ, or (B) IFN-induced Mφ_IFN_ macrophages. For each protein, the fold change was calculated based on their abundance (iBAQs) in V_1_, V_0.5_, or V_0.1_ capsids compared to D capsids using a linear scale from violet being the lowest to dark green being the highest.

**Supplementary Figure S3B:**
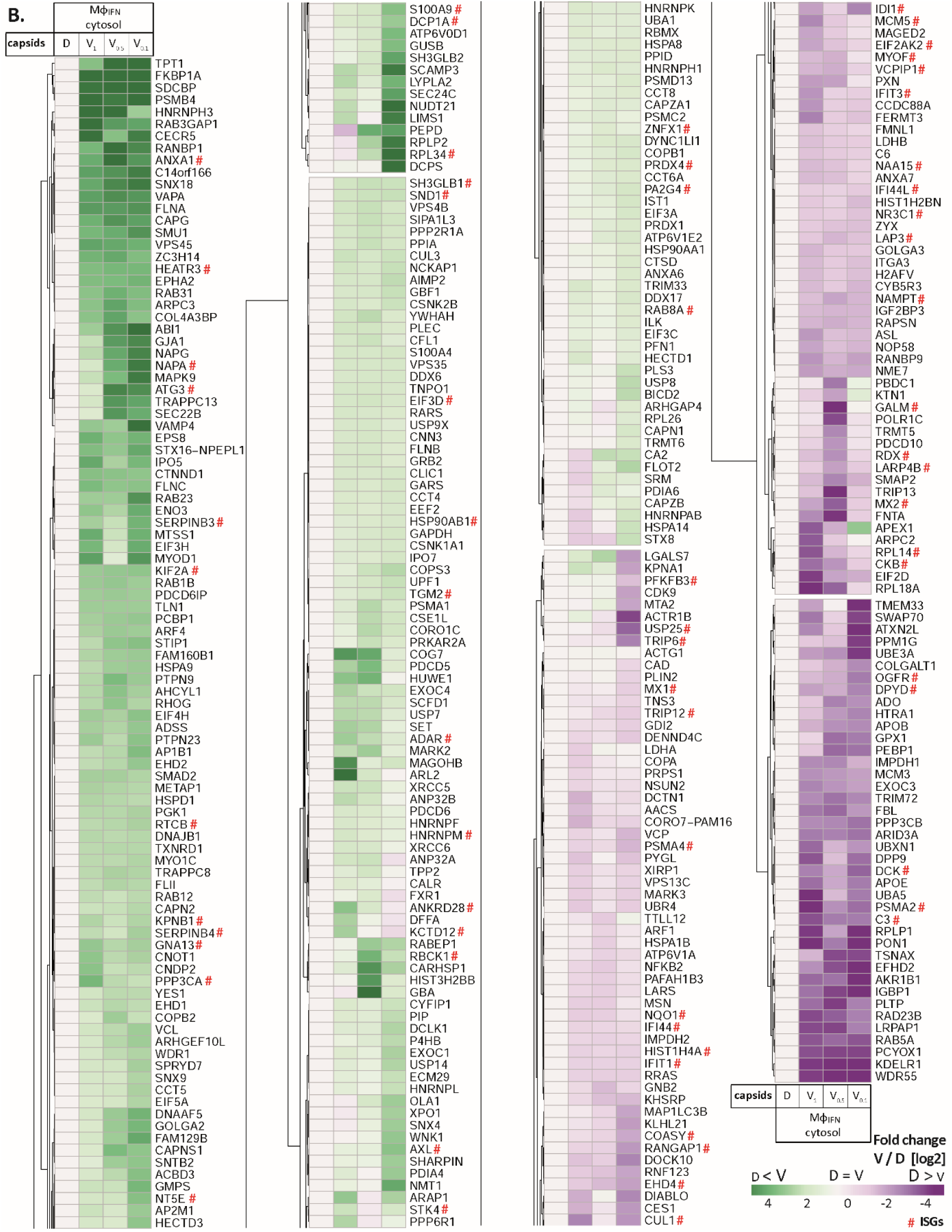
HSV-1 capsids interactomes. Unbiased hierarchical clustered heat map showing the log_2_ fold changes of host proteins identified from capsids-host protein sediments (c.f. Fig. 2; abundance log_2_ difference larger than 1; significance permutation-based FDR smaller than 0.05) from (A) cytosol of resting macrophages (M_φ_), or (B) IFN-induced macrophages (M_φIFN_). For each protein, the fold change was calculated based on their abundance (iBAQs) in V_1_, V_0.5_, or V_0.1_ capsids compared to D capsids using a linear scale from violet being the lowest to dark green being the highest.

**Supplementary Figure S4:**
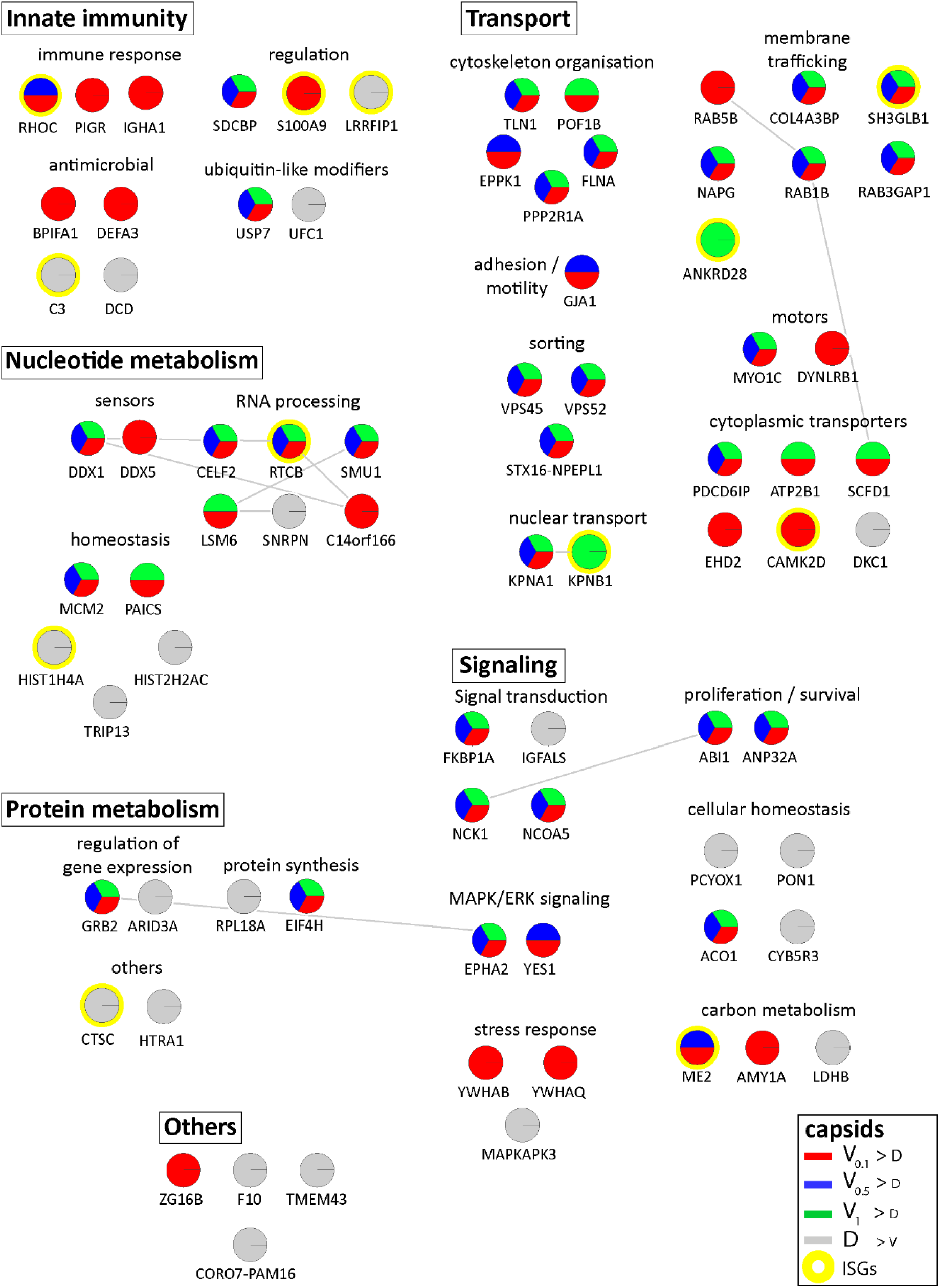
Cytosolic proteins of resting macrophage binding to HSV-1 capsids. *Host* proteins from cytosol of resting Mφ (c.f. Fig. 3A, 3B, 3C; abundance log_2_ difference larger than 1.5; significance permutation-based FDR smaller than 0.05) interacting with V_0.1_, V_0.5_, V_1_, or D capsids were assembled into a functional interaction network of known protein-protein-interactions (grey lines; STRING database, confidence score of 0.7), and grouped according to their known functions (Gene Ontology, Pathway analysis). The Pie chart for each protein indicates its relative enrichment on V_0.1_ (red), V_0.5_ (blue), V_1_ (green), or D capsids (grey).

**Supplementary Figure S5:**
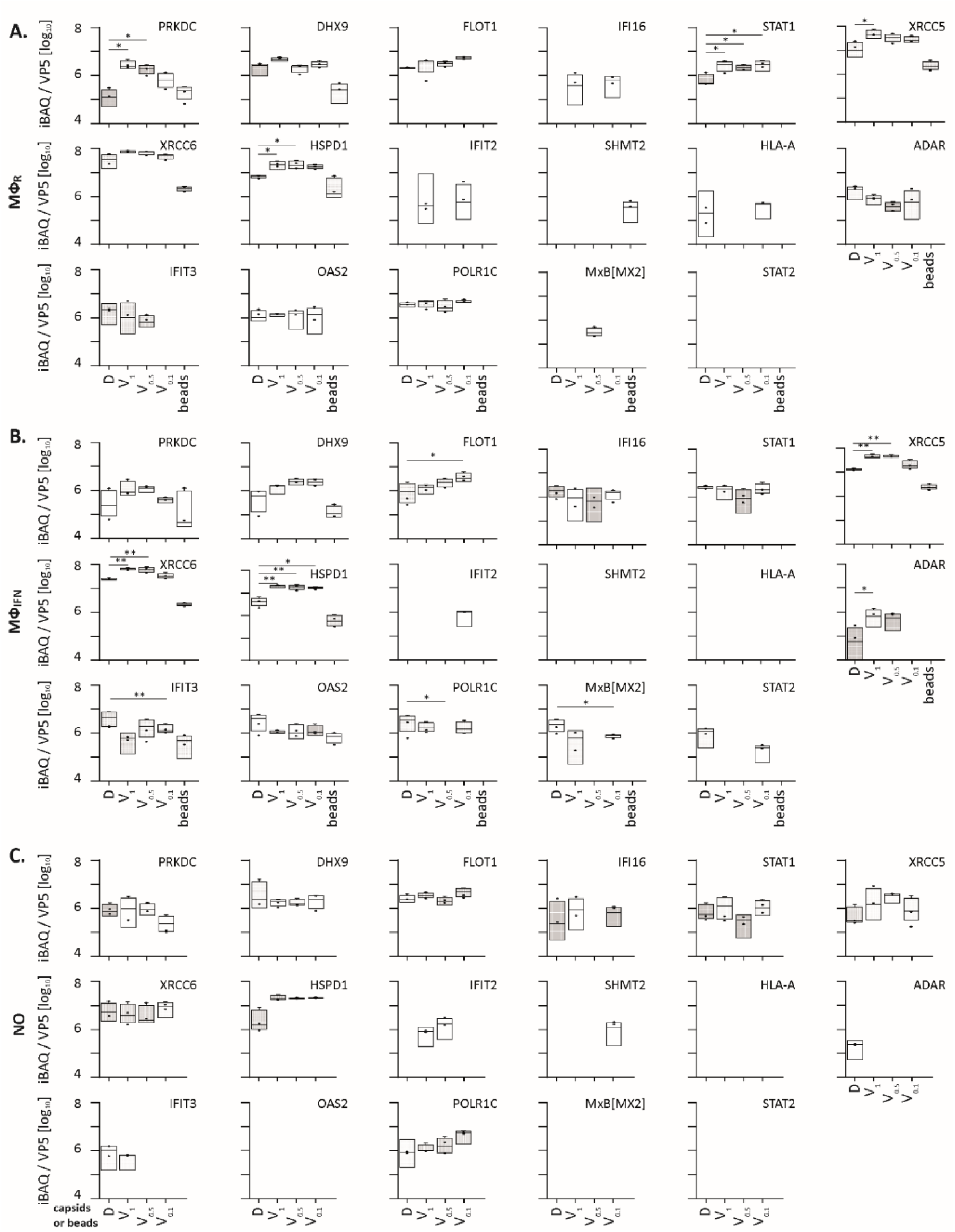
HSV-1 capsids binds to a few ISG proteins. Box and whisker plot of iBAQs showing the differential detection of PRKDC, DHX9, FLOT1, IFI16, STAT1, XRCC5, XRCC6, HSPD1, IFIT2, SHMT2, HLA-A, ADAR, IFIT3, OAS2, POLR1C and MX2 in D, V_1_, V_0.5_ and V_0.1_ capsids-host protein sediments after incubation in (A) cytosol of resting MφR macrophages, (B) IFN-induced MφIFN macrophages or (C) no cytosol. (*) design the significant binding to D or V0.1, V0.5 and V1 capsids as assessed by Welch’s t-test (two-tailed, permutation-based FDR ≤ 0.05) comparing D vs V_0.1_, V_0.5_ or V_1_ capsids in each cytosol separately.

**Supplementary Figure S6:**
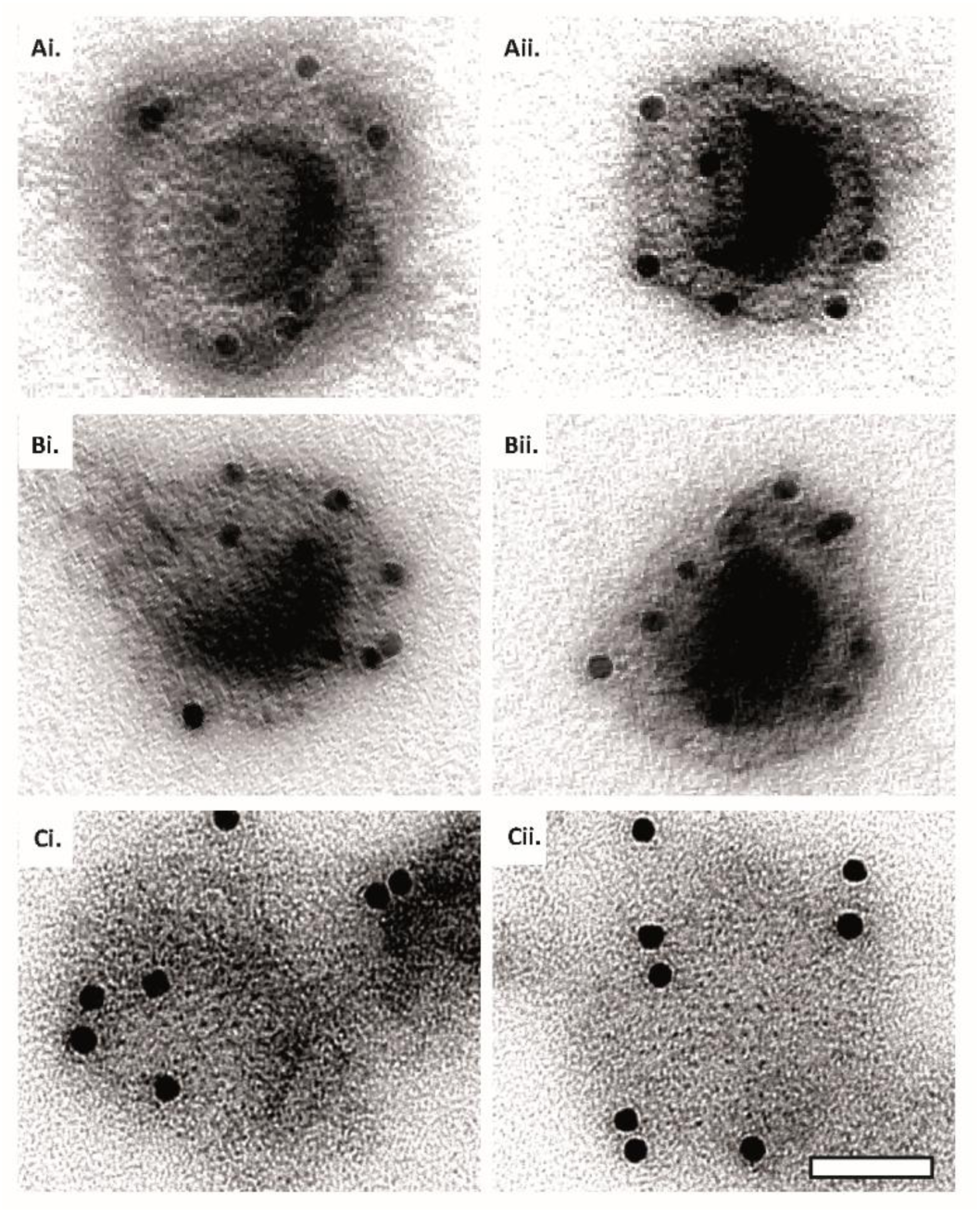
Capsid disassembly intermediates by anti-capsid immunoEM. *Images* of capsids after negative staining and labelling with antibodies raised against the major capsid protein VP5 (NC-1), after incubation in ATP-complemented cytosol from A549 control or MxB(1-715) expressing cells for 60 min at 37°C, and classified as (A) *intact*, (B) *punched*, or (C) *flattened* shells. Scale bar: 50 nm.

**Supplementary Figure S7:**
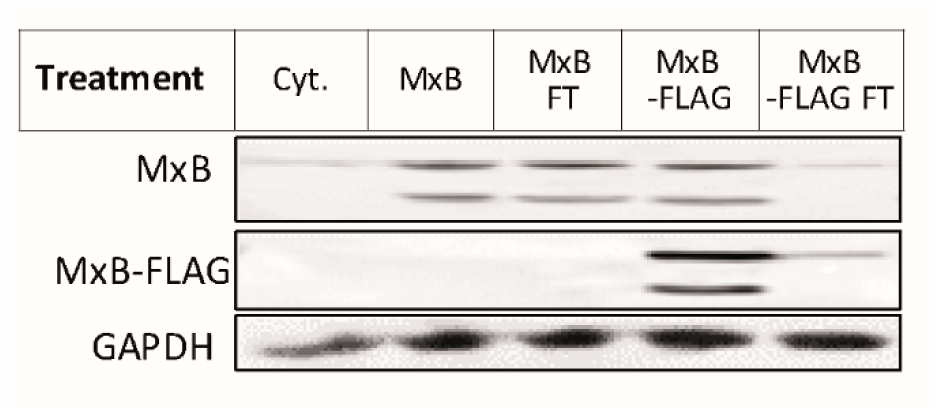
Cytosol immunodepleted for MxB. Cytosols prepared from A549-MxB(1- 715) and MxB(26-715) expressing MxB(1-715) and MxB(26-715), or A549-MxB-FLAG cells expressing MxB(1-715)-FLAG and MxB(26-715)-FLAG, respectively, were incubated with agarose beads coupled to anti-FLAG antibodies. After immunodepletion with anti-FLAG beads to deplete MxB(1-715)-FLAG and MxB(26-715)-FLAG, the flow through (FT) was harvested. To determine to what extend the FLAG- tagged MxB proteins had been depleted, the starting cytosols (MxB, Mxb-FLAG) as well as the respective FT fractions were probed by immunoblot using antibodies directed against MxB, FLAG, or GAPDH as a loading control. **Figure S7-source data 1,2.** Full western blot images for the corresponding detail sections shown in Figure S7.

**Supplementary Figure S8:**
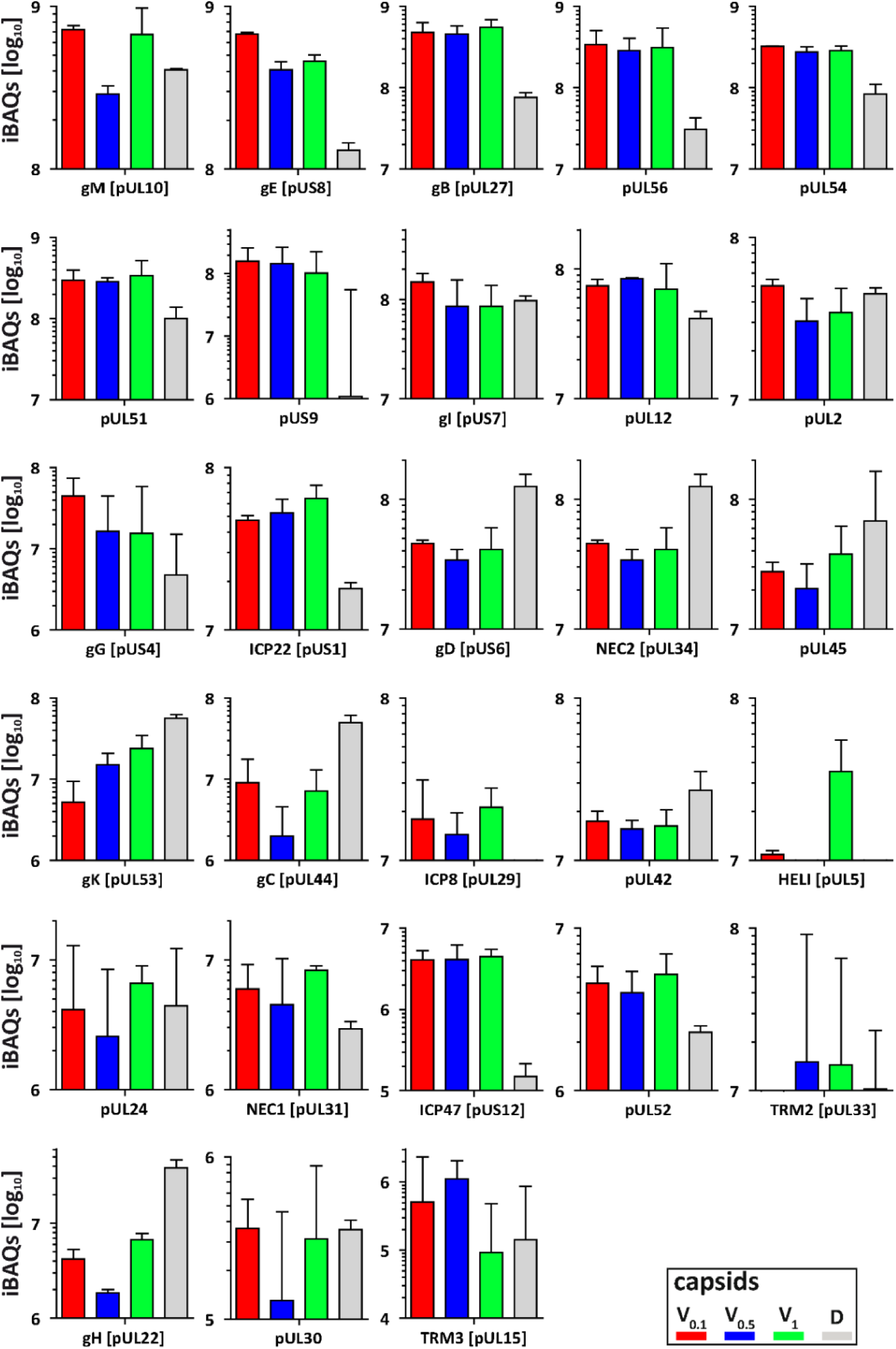
Membrane and non-structural proteins on V capsids versus D capsids. The composition of HSV-1 derived V_0.1_ (red), V_0.5_ (blue), V_1_ (green) and D (grey) capsids were analysed by quantitative mass spectrometry in four biological replicates. The sum of all the peptides intensities (iBAQ, intensity-based absolute quantification) of each viral protein unknown to participate in the structure of the capsids was normalized to the one of VP5 and displayed in a bar plot

